# Three Modes of Viral Adaption by the Heart

**DOI:** 10.1101/2024.03.28.587274

**Authors:** Cameron D. Griffiths, Millie Shah, William Shao, Cheryl A. Borgman, Kevin A. Janes

**Affiliations:** Department of Biomedical Engineering, University of Virginia, Charlottesville, VA 22908, USA; Department of Biochemistry & Molecular Genetics, University of Virginia, Charlottesville, VA 22908, USA

## Abstract

Viruses elicit long-term adaptive responses in the tissues they infect. Understanding viral adaptions in humans is difficult in organs such as the heart, where primary infected material is not routinely collected. In search of asymptomatic infections with accompanying host adaptions, we mined for cardio-pathogenic viruses in the unaligned reads of nearly one thousand human hearts profiled by RNA sequencing. Among virus-positive cases (∼20%), we identified three robust adaptions in the host transcriptome related to inflammatory NFκB signaling and post-transcriptional regulation by the p38–MK2 pathway. The adaptions are not determined by the infecting virus, and they recur in infections of human or animal hearts and cultured cardiomyocytes. Adaptions switch states when NFκB or p38–MK2 are perturbed in cells engineered for chronic infection by the cardio-pathogenic virus, coxsackievirus B3. Stratifying viral responses into reversible adaptions adds a targetable systems-level simplification for infections of the heart and perhaps other organs.

## Introduction

The heart is a cryptic site for viral infections that can be lethal (*1–3*). Patients with viral myocarditis follow a rule of thirds, with roughly equal proportions that i) stabilize, ii) recover spontaneously, or iii) progress in disease severity (*4*). Explanations or hypotheses for these differences are lacking. Cases of viral myocarditis were once diagnosed by endocardial biopsy, but noninvasive imaging is now preferred (*5–7*). This limits molecular analysis to primary heart samples that were either failing or collected for other purposes. Animal surrogates of infection are useful, and the outcome of some cardio-pathogenic viruses differs among mouse strains (*8, 9*); however, they lack the diversity of human populations.

Cardio-pathogenic viruses are frequently detected in hearts failing by dilated cardiomyopathy (*2, 10*), but viral myocarditis can also be asymptomatic (*6, 11*). It is thus unclear what changes and classifications distinguish virus-positive hearts from uninfected ones. One method for summarizing viral landscapes in human tissues (*12, 13*) and cancers (*14–16*) is RNA sequencing (RNA-seq). After aligning raw sequencing data to the human transcriptome, residual unaligned reads are mapped to a suite of candidate pathogens for further study. Viral mining of cancer transcriptomes has shown that premalignant cells adapt and evolve differently when driven by viral oncoproteins (*17*). These results from primary tumors are readily complemented by functional tests in virus-positive cancer cell lines (*18*), but that strategy does not extend easily to other infections. The notion of long-term changes in response to cardio-pathogenic viruses thus remains untested.

Here, we asked whether human hearts organize into a common set of adaptive states after viral challenge. By compiling 979 cardiac transcriptomes from different sources, we identify 189 cases positive for at least one cardio-pathogenic virus. The human transcriptomes of these undiagnosed cases fall into three robust classes, which we consider as “adaptions” to the inciting virus. The adaptions are not specific to any viral species and recur when cells or animals are infected with cardio-pathogenic viruses outside the initial datasets. Follow-up experiments suggest that the adaptions are connected and can switch with specific genetic or pharmacologic perturbations. The triplet of adaptive states sheds light on a convergent response to viral infection of the heart.

## Results

### Various cardio-pathogenic viruses are detected in transcriptomes of the human heart

We mined for undiagnosed heart infections in three deeply sequenced cohorts (**Table 1**) (*19–21*). After filtering the aggregate data based on RNA quality (**fig. S1**), we integrated and analyzed 979 independent samples. Transcriptomes were derived from healthy controls (70.3%) as well as patients with dilated cardiomyopathy (29.7%). More samples were from male donors (64.6%), partly due to the higher prevalence of dilated cardiomyopathy in males (*22*). The inferred genetic ancestry of the samples (*23*) was predominantly European (64.6%), African (17.3%), and Admixed American (16.1%). Together, the aggregate cohort provided a dataset that was clinically and ancestrally diverse.

**Table 1.**
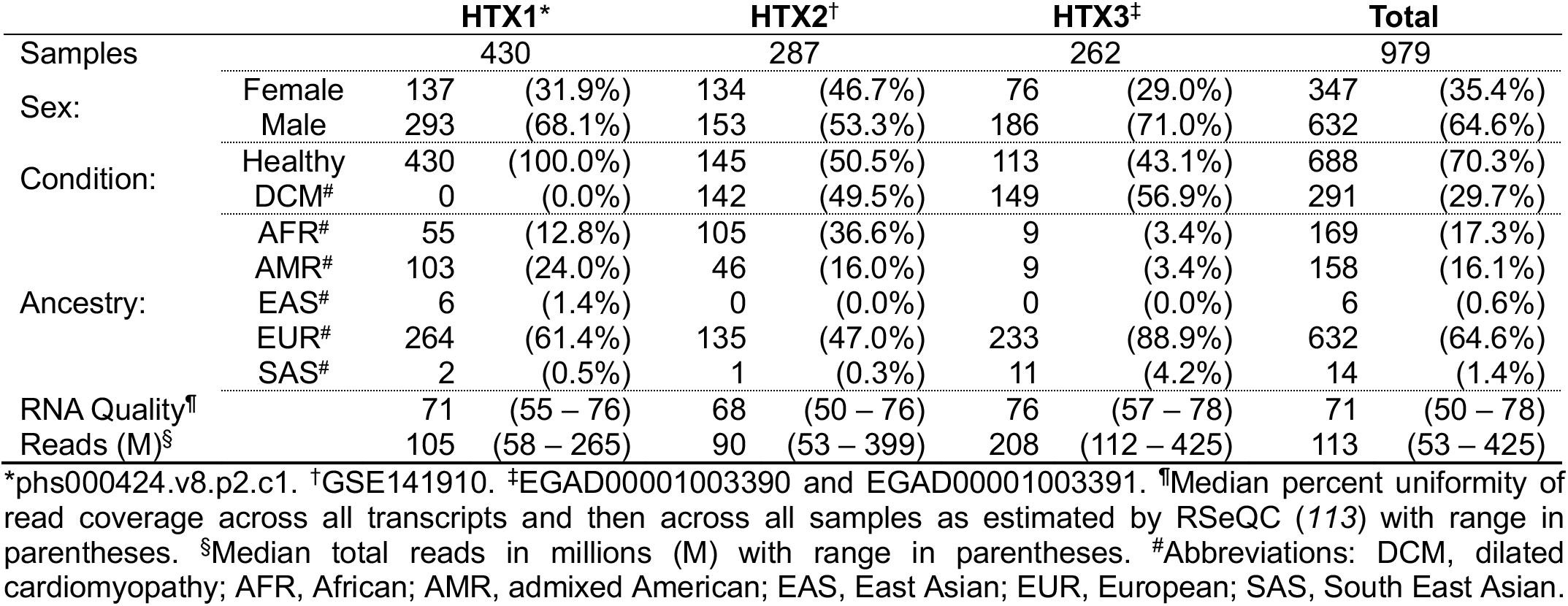
Summary of clinical heart transcriptomics (HTX) datasets used in this study.

Next, we used Virdetect (*24*) to align unmapped reads to cardio-pathogenic viruses previously classified as vasculotropic, lymphotropic, cardiotoxic, or cardiotropic (**Fig. 1A**) (*1*). A total of 189 samples (19.3%) were positive for at least one of nine cardio-pathogenic viruses (**table S1**). In samples with a high read count, we visually confirmed that alignments were consistent with the known regulation of viral transcripts (**Fig. 1A**). Thirteen samples (6.88%) were positive for more than one virus, but there was no positive co-association between any pair of viruses (*P_adj_* > 0.19 by Bonferroni-corrected Fisher’s exact test). The most frequently detected viruses were parvovirus B19, human herpesvirus 4 (Epstein–Barr virus), and adenovirus, consistent with their widespread prevalence (*25, 26*). HIV positivity was an exclusion criteria for HTX1 (*27*), which explains why only one case was detected in this cohort compared to seven cases in HTX2 (both studies were performed in the United States). The elevated presence of human adenovirus C (cardiotropic) in HTX2 is unclear, but geographically clustered cases of adenovirus have been reported in the United States (*28*). Almost all viral transcripts were derived from DNA viruses, indicating active transcription in the heart. In particular, parvovirus B19 transcription distinguishes a clinically relevant infection from a latent bystander (*29, 30*). Thus, asymptomatic virus infection of the heart is not uncommon in humans.

**Fig. 1.**
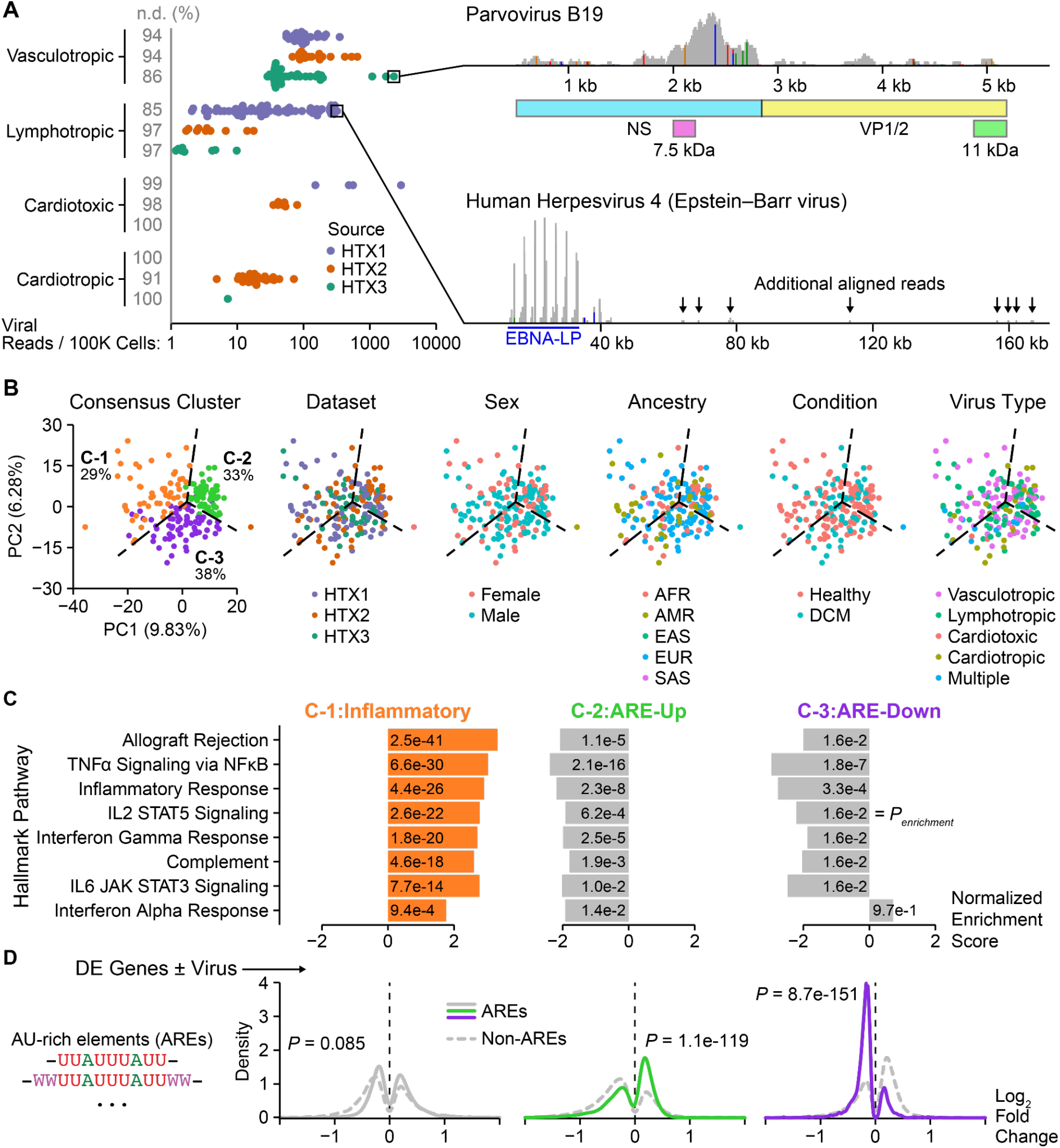
Transcriptomic identification and classification of human heart samples harboring cardio-pathogenic viruses. (**A**) Evidence for vasculotropic, lymphotropic, cardiotoxic, and cardiotropic viruses (*1*) in human transcriptomic (HTX) datasets (Table 1). The percentage of samples in which each class of virus was not detected (n.d.) is shown in gray. Representative alignments to the parvovirus B19 and human herpesvirus 4 genomes are shown above relevant gene products for each virus: NS, 7.5-kDa, and 11-kDa nonstructural proteins, VP1/2 structural proteins, and Epstein-Barr virus nuclear antigen leader protein (EBNA-LP). (**B**) Principal component (PC) projection of the 1000 most-variable genes among virus-positive heart samples (*n* = 189). Samples are colored according to their discrete consensus cluster (fig. S3); HTX source (Table 1); sex; inferred African (AFR), Admixed American (AMR), East Asian (EAS), European (EUR), or South Asian (SAS) ancestry; cases with or without dilated cardiomyopathy (DCM); and type of cardio-pathogenic virus (*1*). (**C**) MSigDB analysis of hallmark pathway enrichments for the three consensus clusters: C-1, Inflammatory; C-2, ARE-Up; and C-3, ARE-Down. The normalized enrichment score is shown for differentially increased (positive) or decreased (negative) genes relative to virus-negative controls. The false discovery rate-corrected *P* value for the enrichment (*P_enrichment_*) is inset. The full set of enrichments is listed in table S4. (**D**) The differentially expressed (DE) genes of C-2 and C-3 show biased proportions of transcripts with AU-rich elements (AREs). The distributions of ARE-containing genes from the ARED-Plus (*129*) database were plotted along with non-ARE DE genes for each cluster, and the DE genes were assessed for ARE enrichment by the hypergeometric test with all genes (DE and non-DE) as the reference and Bonferroni correction for multiple-hypothesis testing.

### Virus-positive hearts cluster into three adaptive responses

To identify commonalities in adaptive responses, we batch corrected the three heart datasets for study, sex, and disease state and performed consensus clustering on the virus-positive samples (**fig. S2**) (*31, 32*). The analysis supported three stable clusters (C-1,2,3) that were comparably sized and not enriched for any available covariate, including virus type (*P_adj_* > 0.95; **Fig. 1B**, **figs. S3** and **S4**, and **table S2**). To identify transcriptional changes characteristic of each virus-positive cluster, we performed differential expression analysis against the virus-negative samples, followed by enrichment analysis (*33, 34*). Among all enrichments, C-1 was strongly and specifically enriched for multiple inflammation-associated hallmarks, including those downstream of NFκB and STAT transcription factors (**Fig. 1C and tables S3** and **S4**). To discriminate between C-2 and C-3, we investigated AU-rich elements (AREs), which are sequence motifs that destabilize transcripts unless there is stress-adaptive signaling from the p38–MK2 pathway (*35*). The differential expression of ARE-containing transcripts robustly separated the two clusters, with highly disproportionate increases in C-2 and decreases in C-3 (**Fig. 1D**). Based on the enrichments, we renamed the consensus clusters *Inflammatory* (C-1), *ARE-Up* (C-2), and *ARE-Down* (C-3). The stratification of virus-positive HTX samples into distinct molecular groups suggested three heart adaptions to infection shared by multiple viruses.

Since the gene-expression differences between clusters could be related to changes in the composition or state of constituent cells in the heart, we used CIBERSORTx (*36*) to estimate abundances of major heart cell types (**Fig. 2A and table S5**). The C-1:Inflammatory and C-3:ARE-Down clusters had somewhat lower abundances of cardiomyocytes compared to virus-negative controls, but C-2:ARE-Up cases were decidedly higher. This phenotype is characteristic of the cardiac hypertrophy that can occur after cardio-pathogen infection (*37, 38*). The C-1:Inflammatory cluster was enriched for fibroblasts, which is consistent with cardio-pathogenic viral infections causing heart fibrosis (*39, 40*). Fibroblasts and endothelial cells were both depleted from the C-2:ARE-Up cluster, possibly due to cell death from infection or a decreased relative abundance compared to cardiomyocytes that had expanded (*40–43*). Corroborating its gene signature, the C-1:Inflammatory cluster showed an increased composition of myeloid and lymphoid cells. Influx of activated innate and adaptive immune cells is synonymous with myocarditis (*1*).

**Fig. 2.**
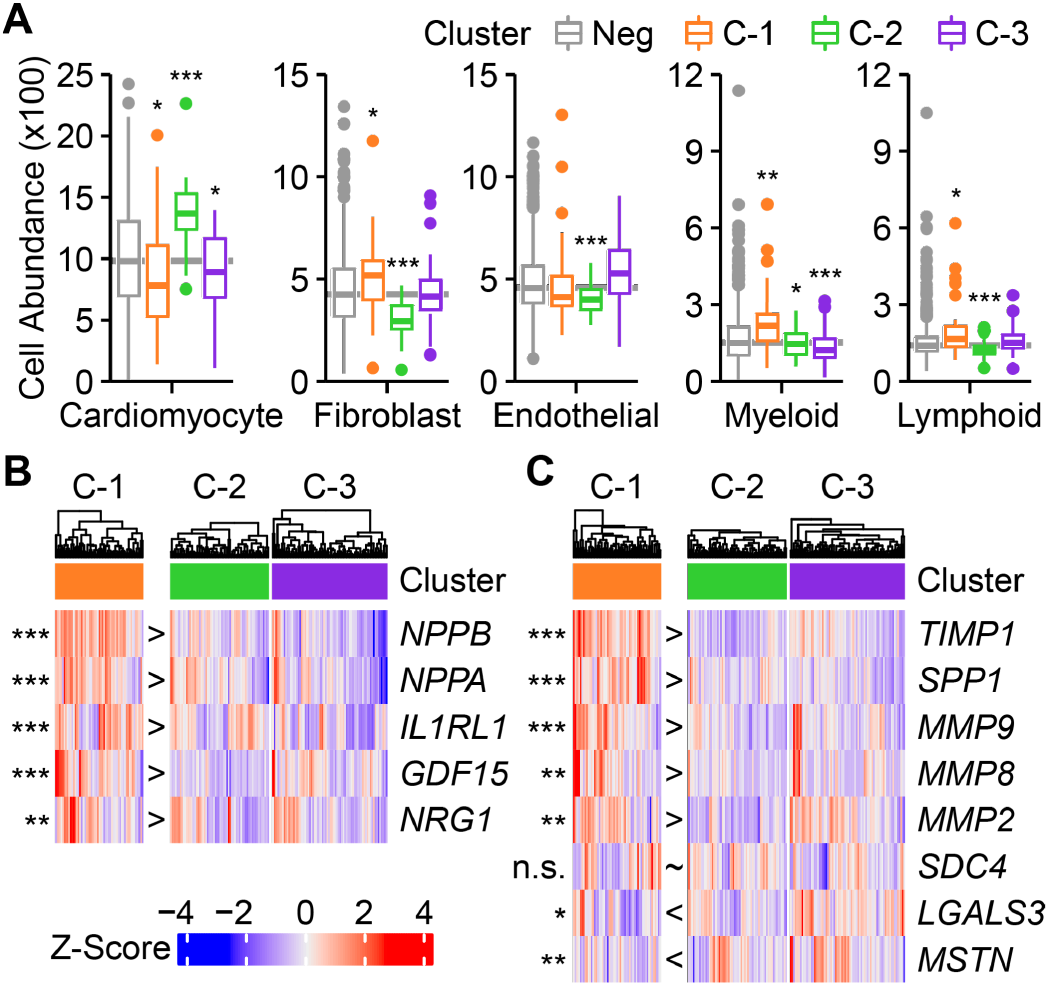
Cellular and molecular phenotypes of the three viral adaptions. (**A**) Deconvolution of bulk heart samples into constituent cell types with CIBERSORTx (*36*). The signature matrix (table S5) was created from a left ventricle single-nucleus RNA-seq atlas of the heart (*121*). Differences in absolute cell type composition between a virus-positive consensus cluster (C-1:Inflammatory, C-2:ARE-Up, C-3:ARE-Down) and the virus-negative reference (Neg) were assessed by multiple *t* tests with the Benjamini-Hochberg correction for multiple hypothesis testing. (**B**) The C-1:Inflammatory cluster is elevated for transcriptional biomarkers of heart failure (*45*). Z-scores of DESeq2-normalized transcript counts are shown for each heart sample, divided by cluster. (**C**) The C-1:Inflammatory cluster shows altered abundances of genes involved in matrix remodeling (*45*). For (B) and (C), differences between C-1 and C-2,3 were assessed by multiple t-tests with the Benjamini-Hochberg correction for multiple hypothesis testing. \**P* < 0.05, \*\**P* < 0.01, \*\*\**P* < 0.001.

Viral heart infections are associated with progression to heart failure (*1–3, 44*), prompting an examination of recognized circulating biomarkers (*45*). Transcripts for multiple heart-failure biomarkers were consistently elevated in C-1:Inflammatory cases compared to the other two clusters (**Fig. 2B**), reinforcing the link between inflammation and heart failure (*46*). Viral heart infections also trigger fibrosis and extracellular matrix remodeling tied to cardiomyopathy (*47–49*). As expected from the increase in fibroblasts (**Fig. 2A**), the C-1:Inflammatory cluster was specifically increased for 5/8 biomarkers of matrix remodeling (*45*) (**Fig. 2C**). These molecular signatures—together with the exchange of cardiomyocytes for fibroblasts and immune cells (**Fig. 2A**)—imply that cases in the C-1:Inflammatory state are actively remodeling and thus unstable. Conversely, the reciprocal patterns for C-2:ARE-Up suggest that it may be stable and physiologically adaptive, with C-3:ARE-Down lying in between.

### Viral adaptions recur *in vitro*

The Inflammatory, ARE-Up, and ARE-Down clusters suggested that adaptions did not depend on underlying characteristics of the heart or the virus (**Fig. 1B**). To test the generality of this finding, we engineered a chronic viral infection *in vitro* by using AC16 cardiomyocytes (*50*) and the prototypical cardio-pathogen, coxsackievirus B3 [CVB3 (*51*)]. CVB3 is normally a lytic virus, but there is precedent for 5’ mutations *in vivo* that disrupt the cytopathic effect (*52, 53*), and released infectious particles are not observed during chronic CVB3 infection (*54*). Inspecting the four N-terminal structural proteins (VP1–4) of the mature CVB3 capsid (*55*), we identified a conserved hydrophobic stretch of VP4 that is important to position VP4^N69^ for autocleavage (**Fig. 3, A–C**). We replaced the VP4^A67^ residue (**Fig. 3B**, red) in that stretch with Gly to displace VP4^N69^ from VP2^D11^ (**Fig. 3B**, orange), which has a presumed role in cleavage (*56*). The goal was to block autocleavage without widespread disruptions to folding of the capsid. Conditioned media from transfectants of the CVB3 A67G mutant did not efficiently form viral plaques, confirming defective encapsidation (**Fig. 3D**). In both AC16 cells and induced pluripotent stem cell (iPSC)-derived cardiomyocytes, CVB3 A67G virions complemented with wildtype VP2:VP4 yielded patterns of differential gene expression very similar to wildtype CVB3 (**Fig. 3, E** and **F** and **tables S6** and **S7**). The CVB3 A67G allele thus mimicked chronic cardio-pathogenic infection without lysis.

**Fig. 3.**
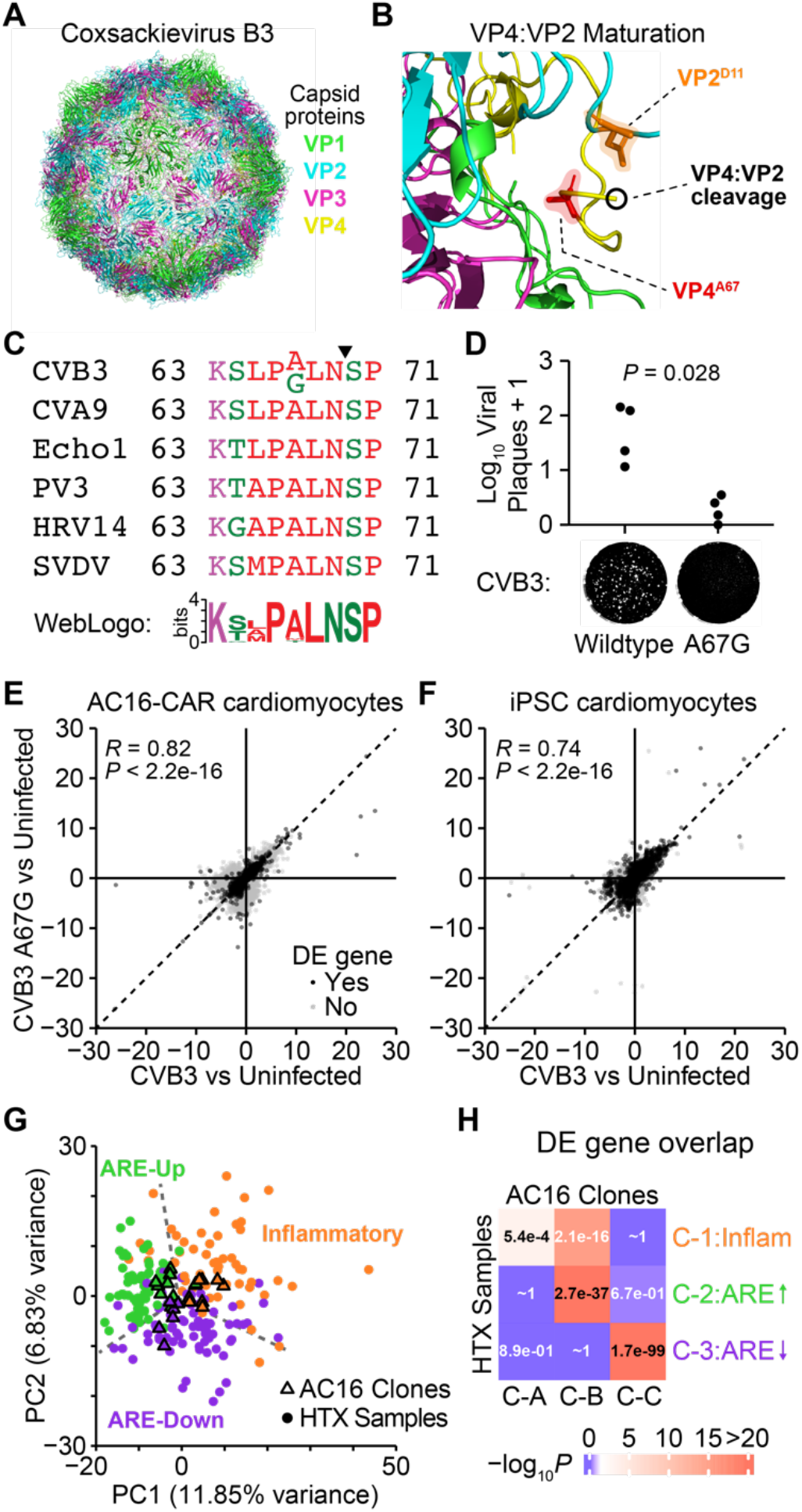
Inflammatory, ARE-Up, and ARE-Down adaptions in an engineered model of chronic cardiomyocyte infection. (**A**) Structural proteins (VP1, VP2, VP3, VP4) of the cardiotoxic virus, coxsackievirus B3 (CVB3). Capsid structure is reprinted from PDB 1COV (*55*). (**B**) Enlarged view of the VP4:VP2 cleavage site of CVB3. The Ala^67^ site of VP4 (VP4^A67^, red) is highlighted along with the putative catalytic Asp11 of VP2 (VP2^D11^, orange). (**C**) Multiple sequence alignment of the residues flanking Ala^67^ among enteroviruses with an N:S cleavage site for VP4:VP2 (arrowhead). The consensus sequence for 30 N:S-containing enteroviruses is shown. (**D**) The CVB3 A67G mutant is not productively infectious. 293T cells were transfected with wildtype CVB3 or the A67G mutant (*n* = 4 biological replicates), and equal titers of released viral RNA were tested for serial infectivity by plaque assay followed by rank-sum test. One replicate of each condition is shown. (**E** and **F**) VP4:VP2-complemented CVB3 A67G infections yield similar transcriptional changes as wildtype CVB3 infections of AC16 cardiomyocytes ectopically expressing CVB3 receptor (AC16-CAR) (E) and induced pluripotent stem cell (iPSC)-derived cardiomyocytes (F). Data are shown as log_2_ fold change, along with the Pearson correlations (*R*) of *n* = 8093 (E) and 9461 (F) differentially expressed (DE) genes. (**G**) Principal component (PC) projection of the 816 genes shared among the 3000 most variable for AC16 clones (triangles) and HTX samples (circles). (**H**) Statistical overlap of DE genes among AC16 consensus clusters (C-A,B,C) and HTX consensus clusters (C-1,2,3).

We stably transfected the CVB3 A67G mutant under the control of a weak promoter (**fig. S5**) into AC16 cells and isolated 12 clonal lines with chronic CVB3 expression to survey adaptive responses. After inducing cardiomyocyte-like quiescence (see Materials and Methods), the 12 clones were transcriptionally profiled and analyzed, yielding three stable consensus clusters (C-A,B,C; **fig. S6**). The three clusters harbored different abundances of CVB3 antisense transcripts without detectable changes in sense transcripts (**fig. S7**), indicating that active viral RNA replication occurs variably among the clones. AC16 is a clonal isolate itself (*50*), raising the further possibility that the observed CVB3 adaptions are partly stochastic.

After intersecting highly variable genes in AC16 clones and human hearts, we noted considerable overlap when the two datasets were combined and projected together (**Fig. 3G**). The leading principal-component scores were smaller for clones, as expected for a single virus-positive cell type in culture, and all were contained within the range of primary samples. We next distinguished each AC16 consensus cluster (C-A,B,C) from the other two by differential gene expression analysis and assessed transcriptome-wide statistical overlaps with the prior heart adaptions (C-1,2,3). Cluster-to-cluster differences were highly concordant between the two datasets (**Fig. 3H**), and follow-up enrichment analyses supported designations of Inflammatory, ARE-Up, and ARE-Down to C-A, C-B, and C-C respectively (**fig. S8** and **tables S8** and **S9**). We conclude that the Inflammatory, ARE-Up, and ARE-Down host adaptions are robust, appearing in two very different infection settings as well as others (see below).

### ARE-Down is an adaptive ground state for the NFκB and p38–MK2 signaling pathways

The AC16 clones provided a context to assess potential relationships among the adaptions. We perturbed the dominant NFκB signature in the Inflammatory adaption by transducing with constitutively active IKK2 [IKK2-EE (*57*)] or IκBα super-repressor [IκBα-SR (*58*); *n* = 3 clones per adaption]. After verifying expression and reciprocal efficacy against direct NFκB targets (**fig. S9, A–C** and **tables S10** and **S11**), we examined collateral changes in other Inflammatory pathway enrichments. IKK2-EE consistently enhanced inflammatory hallmarks in the ARE-Up and ARE-Down groups, but interferon responses for the Inflammatory adaption were dampened (**Fig. 4A** and **tables S12** and **S13**). Chronic interferon signaling from persistent infection may render these cells somewhat refractory to further immune stimulation (*59, 60*). IκBα-SR effects were more uniform across inflammatory hallmarks but variable among groups: the Inflammatory and ARE-Up adaptions were widely inhibited, whereas ARE-Down was mostly unchanged. The IκBα-SR result, together with the low overall expression of NFκB target genes in ARE-Down (**fig. S9B**), indicated that this adaptive state lacked basal NFκB signaling activity.

**Fig. 4.**
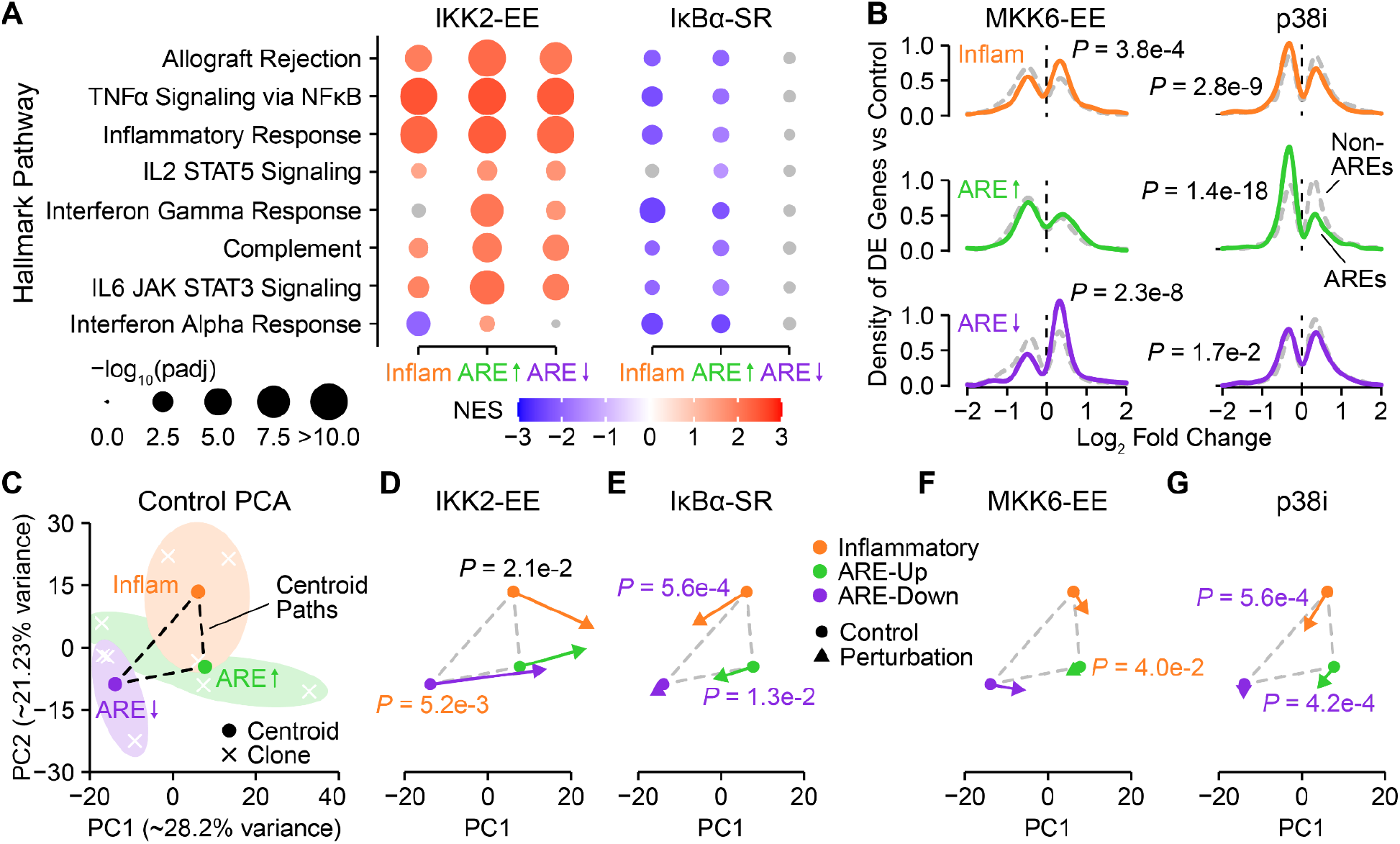
Perturbations of inflammatory and ARE-adapted states. (**A**) Differential changes in inflammatory hallmarks for CVB3-expressing AC16 clones engineered with constitutively active IKK-EE or dominant-negative IκBα super-repressor (IκBα-SR). The normalized enrichment score (NES) was calculated for each inflammatory hallmark pathway relative to empty vector control. The hallmark pathways are ordered as in Fig. 1C. (**B**) Reversion of ARE-Down and ARE-Up states in AC16 clones engineered with constitutively active MKK6-EE or treated with the p38 inhibitor (p38i) BIRB796 (5 µM). The distribution of differentially expressed genes with or without AREs for each adaption was compared as in Fig. 1D. (**C**) Principal component (PC) paths between centroids of adapted states. RNA sequencing data from empty vector controls of each clone are shown and organized by adaption. (**D** and **E**) Displacement of adapted states with IKK-EE, and convergence toward ARE-Down with IκBα-SR. (**F** and **G**) Convergence toward ARE-Down with p38i. Clones (*n* = 3 per adaption) were acutely transduced or treated, quiesced, and profiled by RNA sequencing. For (D) and (E), the relocalization of perturbed centroids (arrowheads) toward the empty vector controls of other states (circles) was assessed by paired two-sided *t* test. Bonferroni-corrected *P* values are colored according to the adaption the centroid moves toward. For IKK-EE, the Inflammatory adaption (black *P* value) moves away from the ARE-Down adaption.

In parallel, we targeted ARE signaling by disrupting the MKK6–p38–MK2 pathway genetically and pharmacologically (*35*). Although transduction of AC16 cells with active MKK6 [MKK6-EE (*61*)] did not lead to constitutive phosphorylation of MK2, we confirmed that ARE-containing genes were preferentially increased (**fig. S9**, **D** and **E** and **table S14**). Conversely, treatment with an allosteric inhibitor of p38 (BIRB796 = p38i) downregulated ARE-containing transcripts, anti-correlating with the gene expression changes elicited by MKK6-EE (**fig. S9, D–F** and **table S15**). Among the adaptions, we noted the greatest impact of MKK6-EE on ARE-Down and of p38i on ARE-Up, as expected, with the Inflammatory group responding to both perturbations (**Fig. 4B**). There was also evidence of adaption crosstalk between the NFκB and p38–MK2 pathways. For example, in the Inflammatory and ARE-Down groups, MKK6-EE strongly inhibited hallmark interferon responses and IKK2-EE differentially increased ARE-containing transcripts, but neither response was observed among clones that were ARE-Up (**fig. S10** and **tables S16** and **S17**). The individual gene–pathway differences across AC16 clones indicated that virally adaptive states could be exaggerated and reset.

To examine the perturbations more holistically, we focused on the centroids of adaptive states and the paths between them along the two leading principal components (**Fig. 4C**). We reasoned that coordinated shifts toward or away from unperturbed centroids would identify changes in adaption at the systems level. By far, IKK2-EE was the most disruptive perturbation, displacing all centroids positively along the first principal component (**Fig. 4D**). Transduced IKK2-EE was expressed very efficiently relative to endogenous IKK2 in AC16 cells (**fig. S9A**), and pure IKK2 holoenzymes are very potent inducers of NFκB (*62*), which likely explains this observation. Nonetheless, the projections remained in the range spanned by virus-positive hearts with the Inflammatory adaption (**fig. S11**), supporting relevance of the IKK2-EE perturbation. More interesting was the impact of IκBα-SR, which caused both the Inflammatory and ARE-Up clusters to move toward ARE-Down (**Fig. 4E** and **fig. S12A).** Even without an overt Inflammatory signature, ARE-Up cells appear to require some level of NFκB signaling to avoid the ARE-Down adaption (**Fig. 4E** and fig. **S8A**). Performing a similar analysis with MKK6-EE, we noted few changes beyond small displacements of the Inflammatory and ARE-Down centroids toward ARE-Up (**Fig. 4F**). By contrast, p38i moved both Inflammatory and ARE-Up adaptions significantly closer to ARE-Down (**Fig. 4G** and **fig. S12B**). Thus, even though the Inflammatory adaption exhibits no differential bias in ARE-containing transcripts, these cells still require p38–MK2 activity to prevent adapting toward an ARE-Down state (**Fig. 4G** and **fig. S8B**). We conclude that the Inflammatory and ARE-Up adaptions are maintained by active signaling from NFκB, p38–MK2 pathways and presumably others. Their absence defines ARE-Down.

### Adaptions are rapid and extend to novel cardio-pathogens

The transcriptomes analyzed thus far were all collected long after challenge with an inciting virus, raising questions about how quickly adaptions emerge. Using wildtype CVB3, we acutely infected iPSC-derived cardiomyocytes and observed significant overlap with the Inflammatory adaption at 8 hours (**Fig. 5**, **A** and **B** and **table S7**). By contrast, CVB3 strongly elicited the ARE-Down adaption when acutely infecting an engineered AC16 derivative with supraphysiological copies of coxsackieviral receptor [AC16-CAR (*63*)] (**Fig. 5**, **C** and **D**, **fig. S13**, and **table S6**). This result implicates ARE-Down in the sequelae of viral infections that overwhelm the host. Together, the *in vitro* studies suggested that some adaptions arise very rapidly.

**Fig. 5.**
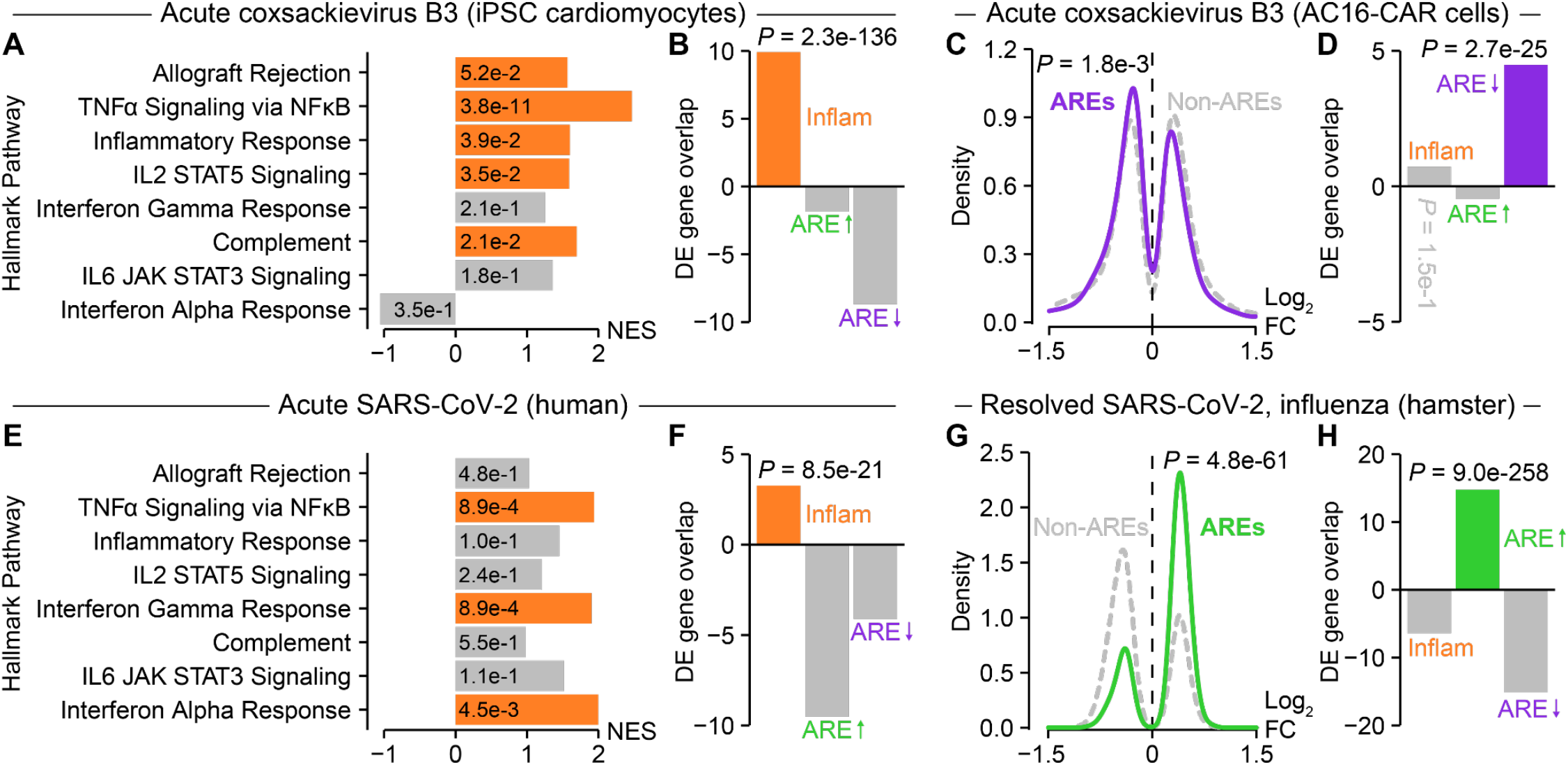
Rapid and divergent viral adaptions to various cardio-pathogens. (**A**) MSigDB analysis of hallmark pathway enrichments among differentially expressed (DE) genes for induced pluripotent stem cell (iPSC)-derived cardiomyocytes infected with coxsackievirus B3 (CVB3) [*n* = 4 samples (control) or 8 samples (wildtype CVB3 MOI = 10 for 8 hours or CVB3 A67G MOI = 3 for 16 hours, each in quadruplicate)]. (**B**) Acute CVB3 infection of iPSC cardiomyocytes triggers the Inflammatory adaption. (**C**) AU-rich element (ARE) analysis of DE genes from AC16 cells overexpressing the CVB3 receptor CXADR (AC16-CAR) infected with CVB3 [*n* = 2 samples (control) or 4 samples (wildtype CVB3 MOI = 10 for 8 hours or CVB3 A67G MOI = 3 for 16 hours, each in duplicate)]. (**D**) Acute CVB3 infection of AC16-CAR cells triggers the ARE-Down adaption. (**E**) MSigDB analysis of hallmark pathway enrichments among DE genes for autopsied hearts from individuals who died from SARS-CoV-2. Data from *n* = 32 cases and 5 controls were obtained from phs002258.v1.p1 (*66*). (**F**) Acute SARS-CoV-2 infection elicits the Inflammatory adaption in human hearts. (**G**) ARE analysis of DE genes from hamsters infected with SARS-CoV–2 or influenza A for 31 days post-infection after histological pathology had resolved. Data from *n* = 5 infections and 3 controls were obtained from GSE203001 (*67*). (**H**) Resolved SARS-CoV-2 and influenza infections elicit the ARE-Up adaption in hamster hearts. For (A) and (E), the hallmark pathways are ordered and displayed as in Fig. 1C by their normalized enrichment score (NES) with false discovery rate-corrected *P* value for the enrichment in the inset. For (B), (D), (F), and (H), the excess overlap of DE genes between the indicated study and the three viral adaptions was calculated and assessed by the hypergeometric test with Bonferroni correction. For (C) and (G), DE gene density is displayed according to log_2_ fold change (FC) and analyzed as in Fig. 1D.

Cardio-pathogen challenge studies are impossible in humans, but surrogates were afforded by recent work with SARS-CoV-2, which can directly infect the heart (*64, 65*). We compared rapid autopsy cases of fulminant SARS-CoV-2 infection (*66*) with uninfected controls and noted the strongest association with the Inflammatory adaption (**Fig. 5**, **E** and **F** and **table S18**). SARS-CoV-2 infects hamsters nonlethally, and we leveraged a recent study (*67*) that profiled the cardiac resolution phase of infection alongside influenza A, a cardio-pathogen we detected earlier in human hearts (**table S1**). We observed the same directional changes in gene expression for both viruses compared to mock-infected controls (**fig. S14**), allowing us to combine them in the analysis. In hamsters, resolution of SARS-CoV-2 and influenza A one month after infection leaves no observable pathology or residual immune infiltration in the heart. We found that resolved hearts showed remarkable overlap with ARE-Up (**Fig. 5**, **G** and **H** and **table S19**), supporting ARE-Up as a favorable long-term adaption to cardio-pathogens. Overall, the recurrence of Inflammatory, ARE-Up, and ARE-Down in vastly different settings leads us to conclude that they represent the three major adaptions for a virus-infected heart.

## Discussion

In this study, we detect cardio-pathogenic viruses in ∼20% of human hearts and find three interconnected adaptions to viral burden that are conceptually organized in **Fig. 6**. The heart normally operates in a state of low NFκB activity (*68*), which increases during acute infection by signaling from pattern recognition receptors for viral nucleic acids (*69*). This Inflammatory adaption is characterized by markers of immune cell influx, matrix remodeling, and heart failure. If NFκB activity is dampened by negative feedback (*70, 71*), the heart progresses to ARE-Up or ARE-Down, depending on p38–MK2 activity (**Fig. 6**, solid lines). The ARE-Up adaption represents resolved infection, consistent with the role of p38–MK2 (*72*). By contrast, the complete lack of p38–MK2 and NFκB signaling in the ARE-Down adaption is more enigmatic and warrants future investigation. It is possible that cells might bypass the Inflammatory adaption entirely or use ARE-Up as an intermediate to reach the ARE-Down state (**Fig. 6**, dashed lines). The p38–MK2 signaling that differentiates ARE-Up from ARE-Down could be a result of ribosomal stress, oxidative stress, or inflammatory cytokines associated with infection and acute cardiac damage (*73*). Viruses specifically activate p38–MK2 during infection (*74*), and they may also impact ARE-containing transcripts independently of p38–MK2. For instance, the 3C protease of CVB3 cleaves an ARE-binding protein to cause an increase in ARE-containing transcripts (*75*). The RNA exosome itself has been reported to antagonize some viruses and promote others (*76, 77*). ARE-Down may mark a state of exhaustion that indicates when a heart is no longer able to actively mount an adaptive response.

**Fig. 6.**
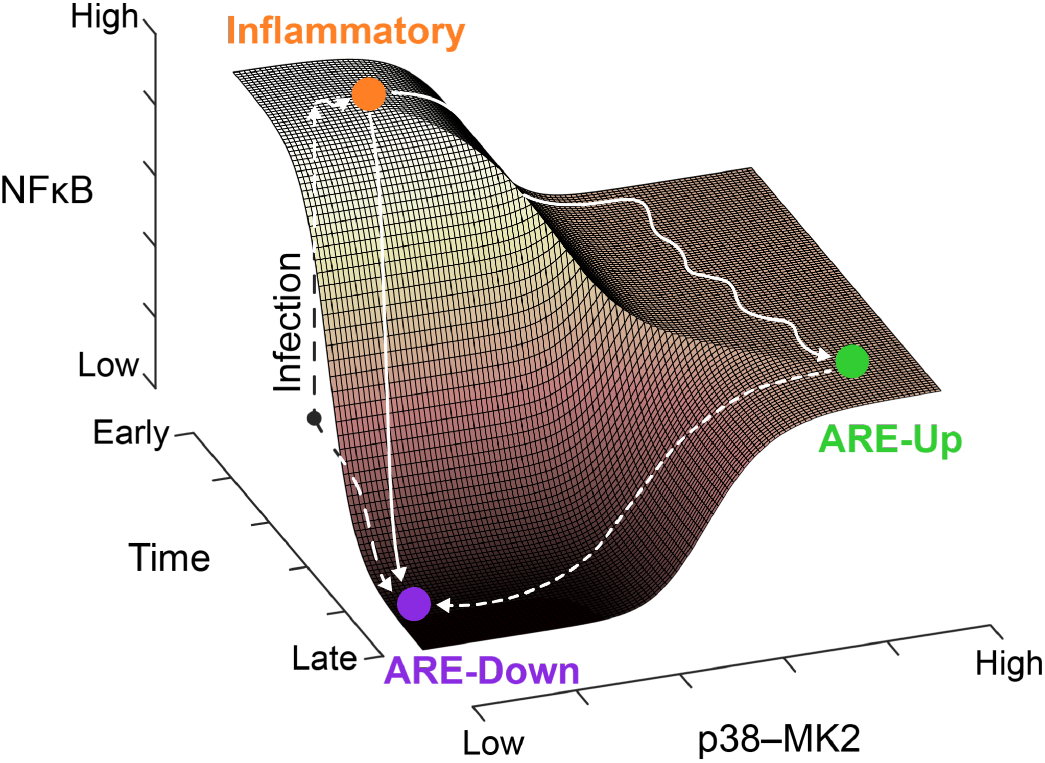
Proposed relationship between heart adaptions after infection. NFκB is acutely activated after infection (dashed vertical), leading to an Inflammatory state that may evolve over time to either the ARE-Up or ARE-Down adaption depending on the trajectory of p38–MK2 signaling (solid). It is unclear whether ARE-Down proceeds through an ARE-Up intermediate (dashed horizontal), and ARE-Down might bypass an Inflammatory intermediate (dashed vertical).

We conceptualize the Inflammatory, ARE-Up, and ARE-Down adaptions along NFκB and p38–MK2 “axes” (**Fig. 6**), but clearly there is crosstalk. In AC16 cells with CVB3, acutely activating or inhibiting p38–MK2 causes NFκB activity to change in the same direction (**fig. S10A**), possibly by impacting IκBα abundance (*78*) or NFκB recruitment via histone phosphorylation (*79*). However, the stable ARE-Up adaption was not enriched for NFκB hallmarks (**Fig. 1C** and **fig. S8A**), suggesting that the two pathways can uncouple. Besides p38–MK2 and NFκB crosstalk, we also noted complex dependencies with the interferon response. Like NFκB target genes (*80*), type I interferon-stimulated genes (*81*) are highly enriched in AREs (*P* < 10^-9^ by hypergeometric test). In ARE-Up cells, interferon hallmarks track with NFκB hallmarks when the NFκB pathway is perturbed (**Fig. 4A**), implying a link during acute perturbation. However, for the Inflammatory adaption, interferon hallmarks are reduced upon NFκB activation and inhibition, as well as upon p38–MK2 activation with MKK6-EE (**Fig. 4A** and **fig. S10A**). Interferon hallmarks are reduced even more so by MKK6-EE in ARE-Down cells (**fig. S10A**), suggesting that attenuators of the interferon response must also be regulated by AREs.

Cardiomyocytes have constitutive type I interferon expression, which primes them to initiate a rapid inflammatory response as a primary alarm (*82, 83*). This may explain the secondary responses to NFκB and p38–MK2 perturbation observed in engineered AC16 cells. Notably, interferon hallmarks do not coincide with NFκB when iPSC cardiomyocytes are infected with CVB3 (**Fig. 5A**), hinting that crosstalk with the interferon pathway is not as general as the adaptions defined by NFκB and p38–MK2.

The three adaptions identified here have implications for the link between cardio-pathogenic virus infection and heart failure (*1–3, 44*). Treatment of active heart infections with antivirals improves cardiac function and prognosis (*30, 44, 84*). Among the ∼20% of people we found with unresolved infection, it is unknown whether intervention might benefit some of the adaptions described. We suspect that the Inflammatory adaption is an undesirable end state (**Fig. 6**), given its increased heart-failure biomarkers and the general role of chronic inflammation in heart failure (*46*). Conversely, the stress-adapted ARE-Up state appears the most favorable at the molecular level (**Fig. 5, G** and **H**), which is cautionary for p38 inhibitor treatments of cardio-pathogens, such as SARS-CoV-2 (*85*). Besides heart failure, viral positivity is associated with cardiac graft rejection (*86*), and there are similarities between viral myocarditis and myocarditis caused by immune checkpoint inhibition for cancer therapy (*87, 88*). It will be interesting to examine how virally adaptive states intersect with other cardio-pathologies. Further, transcript abundances of approved therapeutic targets for heart failure differ significantly among states (**fig. S15**), suggesting that they may also impact therapeutic responsiveness.

Although we repeatedly detected nine species of cardio-pathogenic viruses in all four recognized categories (*1*), others were certainly missed. For example, we did not find any evidence of cardio-pathogenic enteroviruses such as CVB3, even though they have been detected in human hearts by methods other than RNA-seq (*53, 89, 90*). We believe that sensitivity is compromised by the long double-stranded RNA intermediate of the enteroviral life cycle and the equal positive-negative strand ratios that characterize persistently infected hearts (*91*). Long double-stranded RNA is not sequenced effectively by standard protocols (*92, 93*), and others have been unable to detect enteroviruses in any GTEx sample (*12*). A large meta-analysis of 17,000 samples found CVB3 in only 14 samples, six of which were likely alignment errors because these six samples were experimentally infected with a related enterovirus (*94*). CVB3 infections also occur focally in the heart (*91, 95*), which raises sampling issues when considering the small amount of material needed for RNA-seq. More broadly, viral genomes lacking a poly(A) tail are missed by standard first-strand synthesis protocols unless specialized enrichment or depletion protocols are used (*42, 96–98*). The most readily detected viruses have DNA genomes that persist long-term in human tissue due to integration (*99, 100*), episomal expression (*101*), interferon-damped replication (*102*), or other mechanisms (*103*). Latent DNA viruses can also be detected by whole genome sequencing (*16*), but RNA-seq indicates active transcription from viral loci. Encouragingly, even for SARS-CoV-2 (an RNA virus that emerged after the original HTX samples were collected), we observed adaptions concordant with other cardio-pathogenic viruses, supporting their generality.

A limitation of this work and meta-analyses more generally is that confounders cannot be randomized as in a prospective study. We used batch correction to control for known differences (dataset, sex, and disease state) and noted no residual biases in ancestry, but viral species were unequally distributed across datasets. In addition, the timing between virus exposure and sample acquisition is unknown, and there is no clinical or molecular follow-up. The extensions to cultured AC16 cells and iPSC cardiomyocytes address some of these concerns, but they show less gene expression variability (**Fig. 3G**) and lack multiple other cell types in the heart. Given the dramatic differences, it is striking that human heart samples, cultured cells, and an animal model of heart infection converge upon the same three adaptions. An open question is whether the adaptions extend to chronic viral infections in other tissues (*104*) or to other microbial infections of the heart (*105*). Regardless, the three adaptions described here provide a simplifying framework for interpreting the host response to cardio-pathogenic viruses.

## Materials and Methods

### Plasmids

pcDNA3 CVB3 A67G was prepared by site-directed mutagenesis (Agilent, 200522) of pcDNA3 CVB3 (*106*). pcDNA3 VP4:VP2 (Addgene, 216787) was prepared by PCR cloning of amino acids 1–332 from pcDNA3 CVB3 into the BamHI–EcoRI sites of pcDNA3. pEN_TT CVB3 A67G (Addgene, 216788) was prepared in a two-step procedure: i) PCR cloning of CVB3 from its internal EcoRI site (nucleotides 2764–2769) to the 3’ end (nucleotide 7399), digesting with EcoRI–MfeI, and ligating into pEN_TTmcs (Addgene, 25755) (*107*) digested with EcoRI; ii) PCR cloning of CVB3 from the 5’ end (nucleotide 1) to the internal EcoRI site, digesting with EcoRI, and ligating into intermediate plasmid (i) digested with EcoRI. pSLIK CVB3 A67G hygro was prepared by Gateway recombination of pEN_TT CVB3 A67G (Addgene, 216788) and pSLIK hygro (Addgene, 25737) (*107*) with LR Clonase II (Thermo Fisher, 56484). pcDNA3 CVB3, pcDNA3 CVB3 A67G, and pSLIK CVB3 A67G hygro are BSL2 plasmids that cannot be distributed by Addgene and therefore are available upon request.

pBABE FLAG-IKK2-EE puro (Addgene, 216790) was created by PCR cloning of IKK2-EE (Addgene, 11105) with BclI–MfeI ends and inserting into pBABE puro (Addgene, 1764) digested with BamHI–EcoRI. pBABE 3xHA-MKK6-EE puro (Addgene, 216791) was created by PCR cloning of 3xHA-MKK6-EE (Addgene, 47579) with BglII–SalI ends and inserting into pBABE puro (Addgene, 1764) digested with BamHI–SalI. pBABE IκBα-SR puro was described previously (*108*).

### Cell lines and viruses

#### AC16 and derivatives

AC16 cells (*50*) were originally purchased from Mercy Davidson (Columbia University), and derivation of AC16-CAR cells was described previously (*63*). To isolate clones stably expressing CVB3 A67G, FseI-linearized pSLIK CVB3 A67G hygro was transfected into AC16 cells by using Lipofectamine 2000 (Thermo Fisher, 11668019). Stable integrants were selected with 100 µg/ml hygromycin until control plates had cleared, and clones were selected by limiting dilution into 96-well plates. For experiments, cells were terminally quiesced by shSV40 lentivirus and serum withdrawal as previously described (*106*).

#### AC16 clone perturbations

Amphotropic retroviruses encoding pBABE FLAG-IKK2-EE puro, pBABE 3xHA-MKK6-EE puro, pBABE FLAG-IκBα-SR puro, or empty pBABE puro were produced in HEK-293T cells by calcium phosphate transfection as previously described (*106*). Genetic perturbations were validated by transduction of parental AC16 cells and selection with 2 µg/ml puromycin until control plates had cleared. p38 inhibition was validated by treating parental AC16 cells with 5 µM BIRB796 or 0.05% DMSO for 1 hour, followed by stimulation with 20 ng/ml TNF for 30 minutes. After transduction or treatment, the cells were washed with ice-cold phosphate-buffered saline and then lysed in RIPA buffer [50 mM tris-HCl (pH 7.5), 150 mM NaCl, 1% Triton X-100, 0.5% sodium deoxycholate, 0.1% SDS, 5 mM EDTA supplemented with aprotinin (10 μg/ml), leupeptin (10 μg/ml), pepstatin (1 μg/ml), 1 mM phenylmethylsulfonyl fluoride, microcystin-LR (1 μg/ml), and 200 μM sodium orthovanadate]. Lysates were analyzed by immunoblotting as described previously (*109*).

For perturbation experiments, AC16 clones were transduced with pBABE retrovirus, selected with 2 µg/ml puromycin for 3 days, and replated at equivalent cell density. Transduced AC16 clones were then treated with 5 µM BIRB796 or 0.05% DMSO while undergoing the quiescence protocol (*106*), followed by harvest and RNA-seq. Control cells were transduced with pBABE puro and treated with DMSO to use as a common reference for genetically perturbed cells treated with DMSO and pharmacologically perturbed cells transduced with pBABE puro.

#### iPSC-derived cardiomyocytes

iPSC-derived cardiomyocytes were purchased as an iCell Cardiomyocytes^2^ Kit (Cellular Dynamics, R1017). Cells were cultured in iCell Cardiomyocytes² Maintenance Medium as recommended by the manufacturer and plated in iCell Cardiomyocytes² Plating Medium for two days (asynchronous beating) or four days (synchronous beating) before infection with CVB3. Results from two- and four-day samples were combined in subsequent analyses.

#### CVB3

The Kandolf strain of CVB3 was originally obtained from Bruce McManus (University of British Columbia). Unconcentrated viral stocks were prepared and titered in permissive HeLa cells. VP4:VP2-complemented CVB3 A67G was prepared by transfecting HEK-293T cells with equal molar ratios of pcDNA CVB3 A67G and pcDNA3 VP4:VP2 using Lipofectamine 2000 (Thermo Fisher, 11668019). Viral supernatants were concentrated by ultracentrifugation (100,000 rcf for one hour at 4°C) and titered for encapsidated viral genomes by qPCR compared to stocks of live virus. For the comparison in infectivity between wildtype CVB3 and CVB3 A67G, HEK-293T cells were transfected with pcDNA CVB3 or pcDNA CVB3 A67G using Lipofectamine 3000 (Thermo Fisher, L3000001). After 72 hours, supernatants were collected and half was concentrated by ultracentrifugation (100,000 rcf for one hour at 4°C), followed by RNA extraction using the Qiagen RNeasy Plus Mini Kit (Qiagen, 74134) and quantification of viral RNA by qPCR compared to stocks of live virus. The unconcentrated stocks were then diluted to equivalent amounts of viral RNA and assessed for infectivity by serial dilution and plaque assay as previously described (*106*). For infections of iPSC-derived cardiomyocytes and AC16-CAR cells, wildtype CVB3 was used at an MOI of 10 for 8 hours and CVB3 A67G was used at an MOI of 3 for 16 hours before harvest and RNA-seq.

### Public HTX RNA-seq datasets

#### Alignment

We used three RNA-seq datasets (referred to as HTX1, HTX2, and HTX3) for the human left ventricle. HTX1: 432 samples downloaded from the final (v8) data release of the GTEx Consortium (*19*). HTX2: 166 dilated cardiomyopathy and 166 healthy control samples from the Myocardial Applied Genomics Network (MAGNet) consortium downloaded through the Sequence Read Archive (SRP237337) (*20*). HTX3: 149 dilated cardiomyopathy and 113 healthy control samples downloaded through the European Genome-Phenome Archive (EGA; EGAD00001003390 and EGAD00001003391) (*21*).

HTX1 samples were converted from aligned .bam files to raw .fastq files using samtools v1.12 (*110*). HTX2 samples were converted to raw .fastq files using sratoolkit v2.10.5 (https://hpc.nih.gov/apps/sratoolkit.html#doc). HTX3 samples were converted to raw .fastq files using samtools v1.10/1.12 (*110*) and paired reads were recreated using the fastqCombinePairedEnd.py script from Eric Normandeau (https://github.com/enormandeau/Scripts/blob/master/fastqCombinePairedEnd.py). All samples were examined for read quality using FastQC and MultiQC (*111*) and then aligned with HISAT2 v2.1.0 (*112*) using GRCh38 release 99 as the reference genome and enabling downstream transcriptome assembly option. HTX1-3 datasets were paired-end unstranded.

#### RNA quality estimation

The GRCh38 release 99 .gtf file was converted to a .bed file using UCSC-tools, and the RSeQC Transcript Integrity Number (TIN) package (*113*) was used to estimate RNA quality of each sample based on the uniformity of read coverage for each transcript. Samples with a median TIN < 50 were excluded from analysis (**fig. S1**).

#### Ancestry inference

Genetic ancestry of samples was inferred by calling single nucleotide polymorphisms from RNA-seq reads using freebayes v1.3.4 (*114*) and using Kinship-based INference for Genome-wide association studies (KING) v2.3.0 to classify the samples (*23*). Briefly, the ancestry reference file provided by KING [based on the 1000 Genomes Project and grouped into five super populations (*115*)] was downloaded and converted from GRCh37 to GRCh38 annotations using GATK LiftoverVcf v4.2.3.0, retaining 4,425,241 of 5,768,708 (76.7%) variant sites. The converted reference file was then restricted to variants located in exonic regions, yielding 254,699 variant sites. The GRCh38 reference genome was split into 4420 regions using the split_ref_by_bai_datasize.py script provided by freebayes. Freebayes was then run for each genome region on all samples in HTX1–3, restricting variant calls to those in the KING reference file and filtering for a minimum quality of 20 by vcftools v0.1.16 (*116*). The variant calls for all genome regions were then combined, resulting in 102,233 variant sites called in the RNA-seq samples. The variant sites were input into KING for ancestry inference. When needed, files were converted between PLINK and .vcf formats using PLINK v2.00a20211011 (*117*). Files were indexed, sorted, annotated, concatenated, and exonic regions were extracted using Bcftools v1.9 (*118*).

#### Analysis of cardio-pathogenic virus sequences

Unmapped reads were extracted from HTX1–3 by using samtools v1.12 (*110, 118*). Viral reads were then identified in the unmapped reads by using Virdetect (*24*). Briefly, Virdetect uses the STAR aligner (v2.7.9a) to align unmapped reads to 1893 vertebrate virus reference genomes that have been masked for areas of human homology and low complexity (*24*). Detected viruses were filtered to include only known cardio-pathogenic viruses (*1*) (**table S1**). When samples had reads mapping to multiple viral strains within the same species, the strains were merged into a single species and the highest number of reads was retained. All cardio-pathogenic viruses were inspected using Integrative Genomics Viewer (IGV) (*119*), and those mapping to known plasmid sequences (*13*) or aberrantly across the viral genome were considered negative for that virus—using these criteria, 10 samples with alignments to human adenovirus C and 8 samples with alignments to human herpesvirus 5 were considered negative. Abundance of each virus was calculated as viral reads per 100,000 cells, normalized to the size of the viral genome, the size of the human genome, and the number of human reads in the sample by using the following formula (*120*): 100,000 x [2 x (number of viral reads/viral genome size)]/(number of human reads/size of human exome). Viral species were grouped into vasculotropic, lymphotropic, cardiotoxic, or cardiotropic viral classes (*1*) (**table S1**).

#### CIBERSORTx

We downloaded a single-cell RNA-seq atlas of the human heart (*121*) and subsetted the 486,134 cells to include only single-nucleus RNA-seq of the left ventricle as cardiomyocytes are too large to be observed by conventional single-cell RNA-seq. Doublets and unassigned cells were removed, and the remaining 82,527 cells were randomly subsampled to collect 50 cells (if possible) from each cell type for each donor. The remaining 5036 cells were used to create a signature matrix (**table S5**) for CIBERSORTx (*36*), with quantile normalization disabled, min expression set to zero, five replicates, 0.5 sampling fraction, kappa value of 999, q-value of 0.01, minimum 300 and maximum 500 barcode genes per cell type, and filtering non-hematopoietic genes disabled. CIBERSORTx was run on the DESeq2 normalized counts from HTX1–3 (after batch correction), using the custom heart atlas signature matrix, S-mode batch correction, disabled quantile normalization, absolute mode, and 100 permutations. Initial analysis was performed with all cell types; however, pericytes were excluded from the analysis since they correlated as expected with endothelial cells. Adipocytes, mesothelial cells, neuronal cells, and smooth muscle cells were excluded because median cell type abundances were low (2–75) compared to cardiomyocytes, fibroblasts, endothelial cells, myeloid cells, and lymphoid cells (≥ 150).

### AC16 cell and iPSC-derived cardiomyocyte RNA-seq

Quiescent AC16 clones, acutely infected quiescent AC16-CAR cells, and acutely infected iPSC-derived cardiomyocytes were lysed with buffer RLT and mRNA extracted with the RNeasy Mini Kit (Qiagen, 74104). Sequencing libraries were prepared with the TruSeq stranded mRNA library prep kit per manufacturer’s recommendations (Illumina, RS-122-2101). Libraries were then pooled and sequenced with a NextSeq 500/550 Mid Output v2 kit (150 cycles) according to the manufacturer’s recommendation (Illumina, FC-404-2001). Pooled samples were run in triplicate for a minimum sequencing depth of 20 million reads per sample. Uninfected AC16-CAR cell RNA-seq data (*106*) were previously deposited and were downloaded from GEO (GSE155312).

For quiescent AC16 clones subject to genetic or pharmacological perturbations, cells were lysed with buffer RLT and mRNA extracted with the RNeasy Plus Mini Kit (Qiagen, 74134). Strand-specific sequencing libraries, after poly(A) pulldown, were prepared by Novogene. Sample quality control was performed using an Agilent 5400 bioanalyzer and samples were sequenced on an Illumina NovaSeq 6000 Sequencing System by Novogene at a minimum depth of 20 million reads per sample.

Output files were examined for read quality using FastQC and MultiQC (*111*) then aligned with HISAT2 v2.1.0 (*112*) using the downstream transcriptome assembly option. AC16 and iPSC-derived cardiomyocytes were paired-end stranded (RF). For AC16 and iPSC-derived cardiomyocytes reference genomes, an artificial chromosome was added to the GRCh38 release 99 reference genome containing open reading frames for the integrated pSLIK CVB3 A67G plasmid: positive and negative strand “genes” for CVB3, neomycin resistance gene, reverse tetracycline-controlled transactivator (rtTa), hygromycin resistance gene, and ampicillin resistance gene. SV40 genes [small t (exon 1 and 2), large T (exon 1 and 2), agnoprotein, VP1, and VP2] were also included in the artificial chromosome because SV40 was involved in creating the AC16 cell line (*50*).

### SARS-CoV-2 autopsy heart RNA-seq

Heart-derived RNA-seq data from 32 SARS-CoV-2 positive autopsies and five healthy control samples (derived from two unique donors) were generously provided by Christopher Mason (data related to GSE169504) (*66*). The samples were examined for read quality using FastQC and MultiQC (*111*) then aligned with HISAT2 v2.1.0 (*112*) using the downstream transcriptome assembly option. Human SARS-CoV-2 autopsy samples were paired-end stranded (RF). For the SARS-CoV-2 autopsy reference genome, an artificial chromosome was added to the GRCh38 release 99 reference genome containing the SARS-COV-2 USA-WA1/2020 (NR-52281) “gene”.

### Hamster heart RNA-seq

#### Alignment

RNA-seq data from eight hamster hearts harvested at 31 days post infection (three SARS-CoV-2, two influenza A, and three mock controls), were downloaded through the Sequence Read Archive (SRP375160) (*67*). Hamster heart samples were converted to raw .fastq files using sratoolkit v2.10.5, then examined for read quality by FastQC and MultiQC (*111*). The samples were aligned with HISAT2 v2.1.0 (*112*) using the downstream transcriptome assembly option. Hamster heart samples were single-end stranded (R). For the hamster heart reference genome, an artificial chromosome was added to the *Mesocricetus auratus* 1.0 genome (downloaded from Ensembl release 106) containing influenza A virus H1N1 isolate A/California/04/2009 (ATCC VR-1805) (segments 1-8) and SARS-COV-2 USA-WA1/2020 (NR-52281) as “genes”.

#### Converting hamster gene names to human orthologs

Hamster gene names were converted to human orthologs using the Ensembl ortholog database, through BiomaRt (*122*). Genes were converted by directly matching gene symbols where possible. When multiple hamster genes mapped to a single human gene, the sum of all mapping genes was taken. When one hamster gene mapped to multiple human genes, the total number of reads was divided between the human genes and rounded to whole numbers. When multiple hamster genes mapped to multiple human genes, the sum of hamster reads were divided between human genes and rounded to whole numbers. As a result, 13,611 out of 15,089 unique mapped hamster genes were converted to human symbols; the rest were excluded from analysis.

### Transcript assembly and counting

Transcripts were assembled and counted with Stringtie v2.1.0 (*123*), restricting assembly to the appropriate reference genome and associated artificial chromosomes. The prepDE.py3 script provided by Stringtie was used to extract count information for downstream differential expression with the following average read lengths: 76 bp (HTX1), 125 bp (HTX2), 100 bp (HTX3), 75 bp (AC16 and iPSC-derived cardiomyocytes), 150 bp (SARS-CoV-2 autopsy and AC16 clones that underwent genetic or pharmacological perturbations), and 151 bp (hamster hearts).

### Clustering and differential gene expression

For HTX1 and HTX2, donor sex was obtained from the accompanying metadata. For HTX3, donor sex was inferred by read counts for *EIF1AY* (≥ 1000 for male) and *XIST* (≥ 10,000 for female). HTX1–3 transcriptomes were batch corrected for study (HTX1, 2, or 3), sex, and disease state (DCM vs. Healthy) by using ComBat-seq (*32*). All samples were normalized with DESeq2 (*33*), followed by a variance-stabilizing transformation (*124*) unblinded to the experimental design. The 1000 most variable genes (determined by greatest variance in transcript abundance) were extracted and analyzed by PCA (with centering and scaling) or UMAP (n_neighbors = 40, spread = 10, and min_dist = 1) (*125*). To find clusters, the top 2–3% of variable genes (top 2% = 1085 for human hearts and top 3% = 1067 for clones, both determined by median absolute deviation) were used for Monte Carlo reference-based consensus clustering (M3C) (*31*) to identify a statistically preferred number of clusters. M3C was used with the following clustering parameters: five Monte Carlo iterations, 10 cluster maximum, 0.8 resampling fraction, reverse-PCA reference method, 100 reference resamplings, 100 real data resamplings, spectral clustering algorithm, 0.1 first pac score coordinate, 0.9 second pac score coordinate, 123 seed, entropy objective function, Monte Carlo simulation method, 0.1 default lambda parameter, and tuning lambda on with a range of 0.02–0.1 (intervals of 0.02) for tuning. The cluster assignments and metadata for HTX1–3 are summarized in **table S2**. The differentially expressed (DE) genes (adjusted *P* value < 0.1) specific to each cluster were identified with DESeq2 (*33*) and the following comparisons: infected vs. uninfected (human SARS-CoV-2 hearts, hamster hearts, iPSC-derived cardiomyocytes, and AC16-CAR cells), virus positive cluster vs. all virus negative (HTX1–3), and one cluster vs. the two other clusters (CVB3-expressing AC16 clones). For the CVB3-expressing AC16 clones that underwent genetic or pharmacological perturbations, two analysis designs were used: perturbation vs. control (perturbation effects on all clones together with one DESeq2 object) or perturbation vs. control (separate comparison for each cluster with a DESeq2 object for each perturbation).

### Gene enrichment and overlap analyses

Before gene set enrichment analysis (*34*), the differential log_2_ fold change of all genes was shrunk by using the ashr algorithm (*126*). Genes were ranked based on this shrunken log_2_ fold change (genes with a base mean expression < 1 normalized counts were removed), and the fgsea package (*127*) was used to determine enrichments in the Hallmark pathway dataset (*128*). For enrichment of ARE-containing genes, a complete list of 4884 genes with an ARE in the 3’ untranslated region was downloaded from ARE-Plus (*129*) and the overlap of DE genes with ARE-containing genes was evaluated by the hypergeometric test. Only protein-coding genes retained by DESeq2 were included in the total number of genes for the hypergeometric test.

Enrichment of ARE-containing genes among interferon stimulated genes was similarly tested. All protein-coding genes upregulated >2 fold by interferon-α treatment in human cells were downloaded from Interferome V2 (*130*) and the overlap with ARE-containing genes was assessed by the hypergeometric test. Overlaps between cluster-or dataset-specific DE genes were also evaluated by the hypergeometric test. Only DE genes that were differential in the same direction were considered overlapping. The total number of genes included in the hypergeometric test was the number not filtered out by DESeq2 in either dataset multiplied by two to account for the possibility of overlapping in either the up or down direction. The excess overlap between other datasets and human heart clusters was calculated by subtracting the expected percent overlap from the observed percent overlap using the following formula: 100 x (number of overlapping genes/number of human heart cluster DE genes) - 100 x [number of other dataset DE genes/(total number of genes x 2)].

### CVB3 structure visualization

The CVB3 capsid structure (1COV) was downloaded from PDB (*55*) and visualized using PyMOL version 2.6 (Schrödinger).

### Enterovirus multiple sequence alignment

UniProt was searched for “enterovirus AND (taxonomy_name:enterovirus)”, restricting entries to reviewed (Swiss-Prot) proteins with a sequence length greater than 801. The resulting 44 enteroviruses were aligned with Clustal Omega (*131*). The alignment was subsetted in Jalview (*132*) to include only the 30 enteroviruses with an N:S cleavage site for VP4:VP2. Aligned amino acids 63–71 were input into weblogo (*133*) to generate a consensus motif.

### Quantitative PCR

CVB3 RNA purified from 1 ml conditioned medium was ethanol precipitated and used for first strand cDNA synthesis as previously described (*134*), substituting 2 pmol CVB3.1R primer (**table S20**) for 25 pmol oligo(dT)_24_. Quantitative PCR was performed on a Bio-Rad CFX96 Touch Real-Time PCR Detection System as previously described (*134*) with CVB3.2F and CVB3.2R primers (**table S20**) and an annealing temperature of 55°C.

### Immunoblotting

Near-infrared fluorescence-based immunoblotting was performed as previously described (*109*) and imaged on a LI-COR Odyssey instrument. Primary antibodies recognizing the following targets were used: FLAG (Sigma, F7425, 1:2000 dilution); HA (Roche, 11867423001, 1:1000 dilution); IKK2 (Cell Signaling Technology, 8943, 1:1000 dilution); IκBα (Cell Signaling Technology, 4814, 1:1000 dilution); MK2 (Assay Design, KAP-MA015, 1:1000 dilution); MK2 p-Thr344 (Cell Signaling Technology, 3007, 1:1000 dilution); p38 (Santa Cruz Biotechnology, sc-535, 1:5000 dilution); p38 p-Thr180/Tyr182 (Cell Signaling Technology, 4511, 1:1000 dilution); tubulin (Abcam, ab89984, 1:20,000 dilution); vinculin (EMD Millipore, 05-386, 1:10,000 dilution).

### Statistical Analysis

Unless otherwise mentioned, all graphical and statistical analyses were done in R (v4.1.3) through RStudio (077589bc, 2021-09-20). The following R packages were used: ashr_2.2-54; biomaRt_2.50.3; circlize_0.4.15; ComplexHeatmap_2.10.0; DESeq2_1.34.0; fgsea_1.20.0; Gen3_0.0.10; ggbeeswarm_0.6.0; jsonlite_1.8.0; M3C_1.16.0; patchwork_1.1.2; RColorBrewer_1.1-2; rstatix_0.7.2; SeuratDisk_0.0.0.9020; Seurat_4.3.0.1; sva_3.42.0; tidyverse_1.3.1; umap_0.2.8.0. All other statistical tests and *P*-value cutoffs are described in the figure captions.

## Acknowledgments

We thank J. Saucerman, M. Civelek, and K. Ellis for reviewing this manuscript, C. Mason, J. Foox, and J. Park for assistance with the SARS-CoV-2 autopsy heart RNA-seq data, E. Farber and S. Onengut-Gumuscu at the Center for Public Health Genomics for performing the RNA-seq (unperturbed AC16 clones, AC16-CAR cells, and iPSCs), M. Sutcliffe for assistance with RNA-seq alignments, and Research Computing at The University of Virginia for providing computational resources and technical support

## Author contributions

Conceptualization, C.D.G., M.S., and K.A.J.; methodology, C.D.G., M.S., and K.A.J.; software, C.D.G.; validation, W.S. and C.A.B.; formal analysis, C.D.G. and K.A.J.; investigation, C.D.G., M.S., W.S., and K.A.J.; resources, C.D.G., M.S., W.S., C.A.B., and K.A.J.; data curation, C.D.G., M.S., and K.A.J.; writing – original draft, C.D.G. and K.A.J.; writing – review & editing, C.D.G., M.S., W.S., C.A.B., and K.A.J.; visualization, C.D.G. and K.A.J.; supervision, C.D.G. and K.A.J.; project administration, C.D.G. and K.A.J.; funding acquisition, C.D.G., M.S., and K.A.J.

## Data and materials availability

RNA-seq datasets have been submitted to the National Center for Biotechnology Information Gene Expression Omnibus database GSE262649.

---

**Fig. S1.**
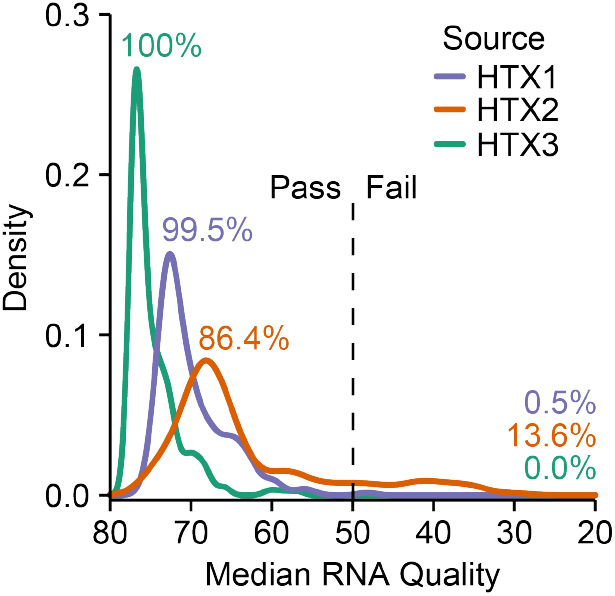
RNA quality assessment of HTX datasets. The median transcript integrity number (TIN) was calculated by RSeQC (*113*) and samples with median TIN < 50 were excluded from further analysis. The percentages of passing and failing samples for each HTX dataset are shown.

**Fig. S2.**
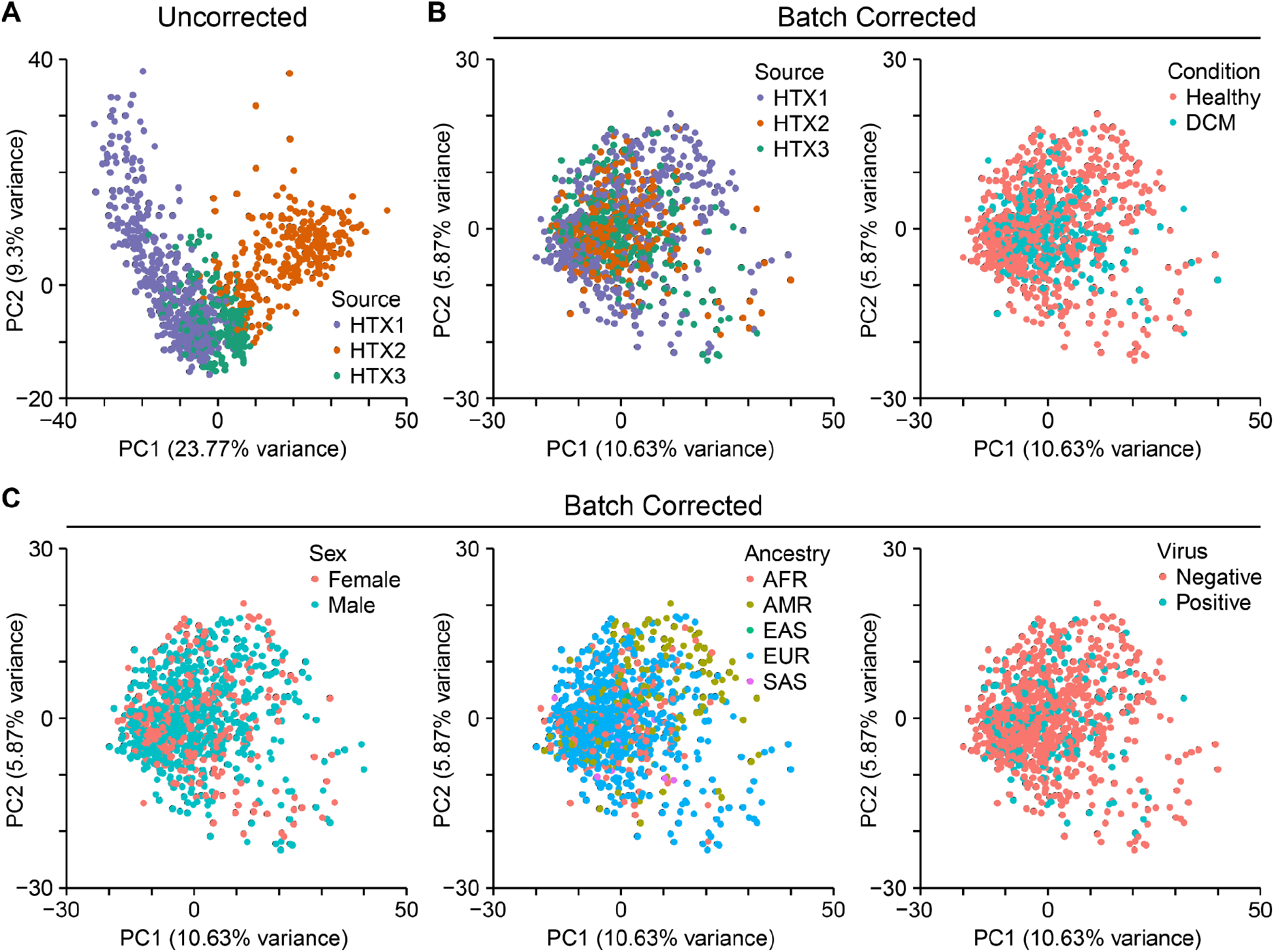
Batch correction of HTX datasets. (**A**) Uncorrected principal component (PC) projection of the 1000 most-variable genes across the HTX datasets (*n* = 979). (**B** and **C**) PC projection after batch correction with ComBat-seq (*32*). Samples are colored according to their HTX source (Table 1); cases with or without dilated cardiomyopathy (DCM); sex; inferred African (AFR), Admixed American (AMR), East Asian (EAS), European (EUR), or South Asian (SAS) ancestry; and positivity for cardio-pathogenic virus (*1*).

**Fig. S3.**
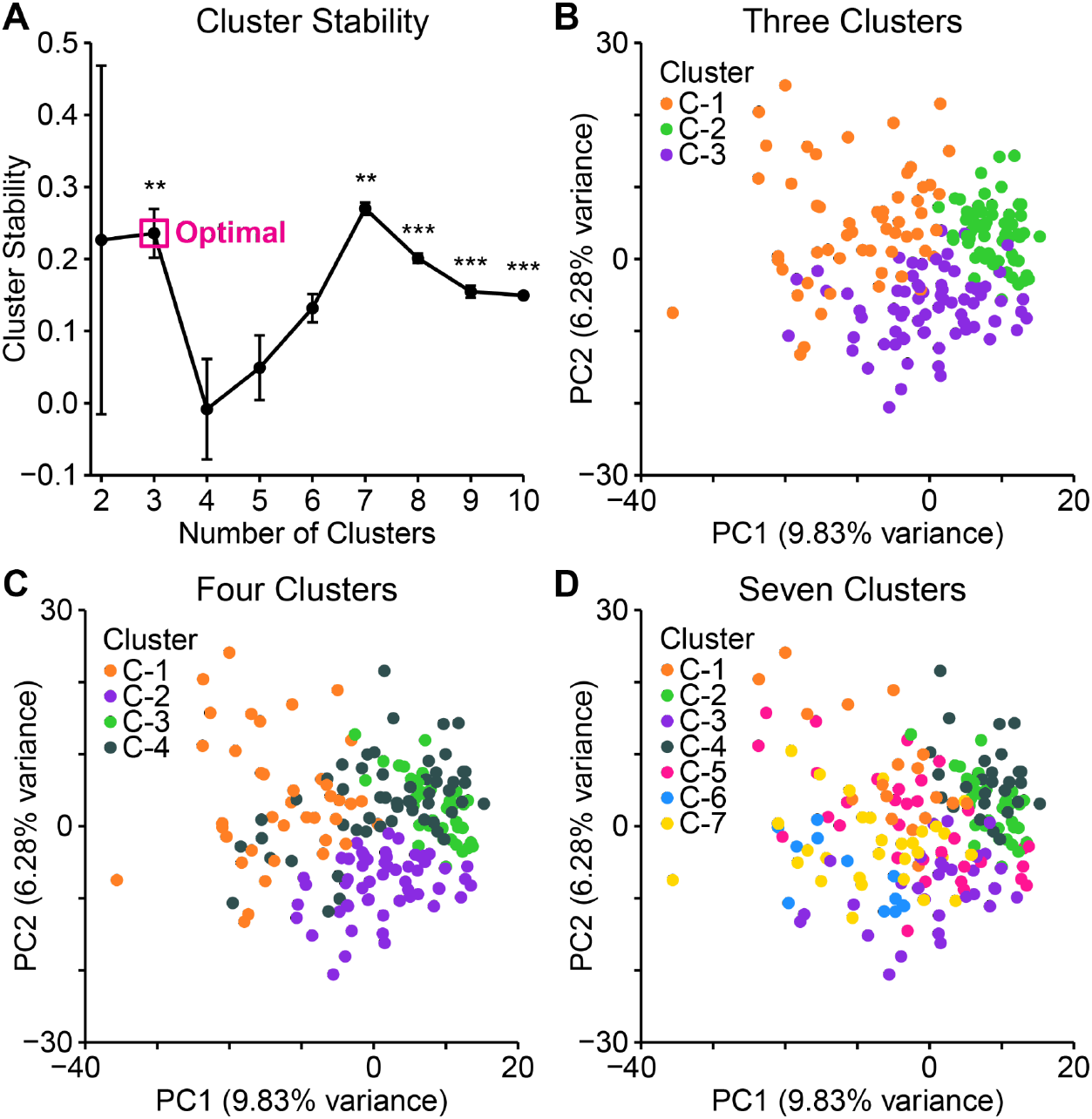
Three consensus clusters of virus-positive heart samples. (**A**) Monte Carlo-based consensus clustering (*31*) stability as a function of the number of clusters. \*\**P* < 0.01, \*\*\**P* < 0.001 for changes in stability between adjacent clusters. (**B** to **D**) Principal component (PC) projection of the 1000 most-variable genes among virus-positive heart samples (*n* = 189). Samples are colored according to their cluster assignment for three, four, and seven consensus clusters.

**Fig. S4.**
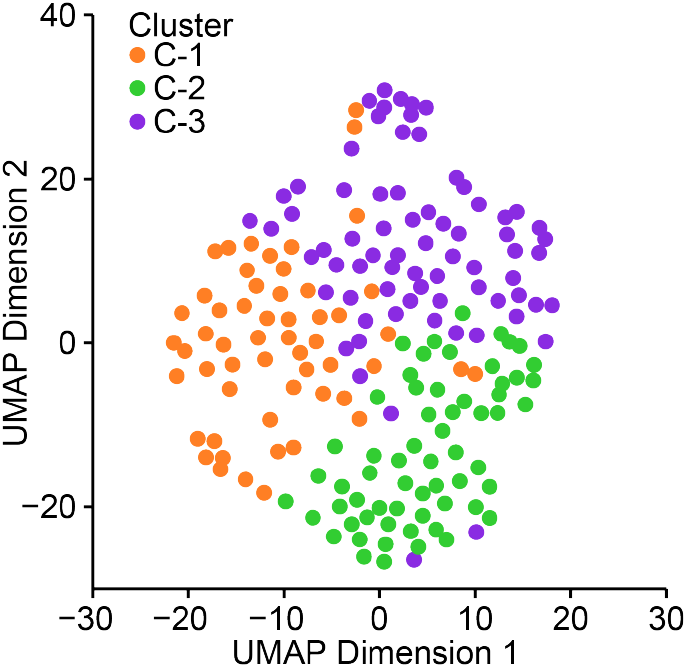
Separability of consensus clusters after nonlinear projection. The 1000 most-variable genes among virus-positive heart samples (*n* = 189) were organized in two dimensions by Uniform Manifold Approximation and Projection (UMAP) (*125*) with 40 nearest neighbors, a spread of 10, and a minimum distance of 1.

**Fig. S5.**
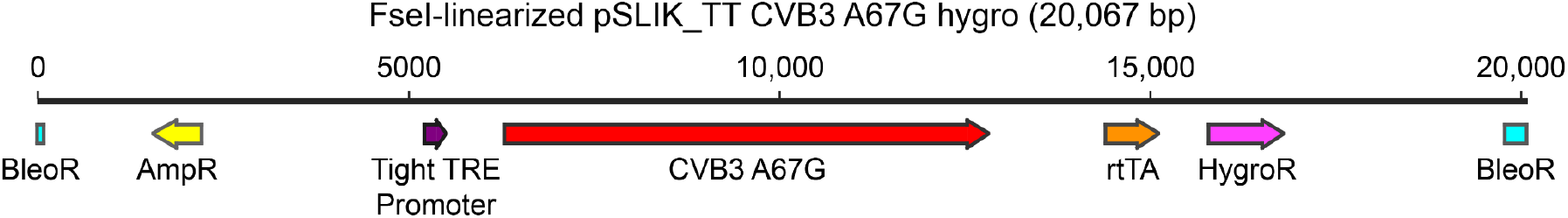
Delivery vector for maturation-deficient coxsackievirus B3 (CVB3). The cDNA for CVB3 A67G was cloned into pEN_TT, recombined into pSLIK hygro, and then linearized with a unique FseI restriction site in the bleomycin resistance (BleoR) cassette. Expression of CVB3 A67G RNA is controlled by a tight tet-responsive element (TRE) and minimal CMV promoter, which weakly drives transcription without doxycycline activation of reverse tet transactivator (rtTA). AC16 cells were transfected with the linearized construct, selected for hygromycin resistance (HygroR), and cloned by limiting dilution.

**Fig. S6.**
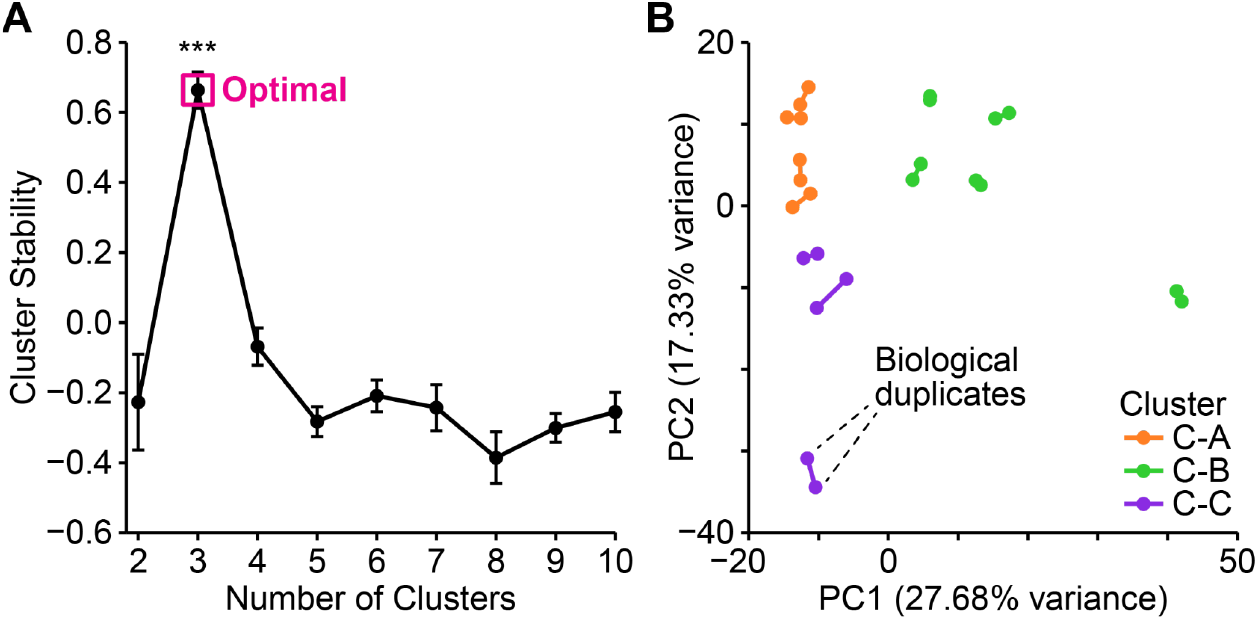
Three consensus clusters of coxsackievirus B3 (CVB3)-positive AC16 clones. (**A**) Monte Carlo-based consensus clustering (*31*) stability as a function of the number of clusters. \*\*\**P* < 0.001 for changes in stability between adjacent clusters. (**B**) Principal component (PC) projection of the 1000 most-variable genes among CVB3-positive clones of AC16 clones (*n* = 12). Samples are colored according to their cluster assignment for three clusters (C-A,B,C), and biological duplicates are paired.

**Fig. S7.**
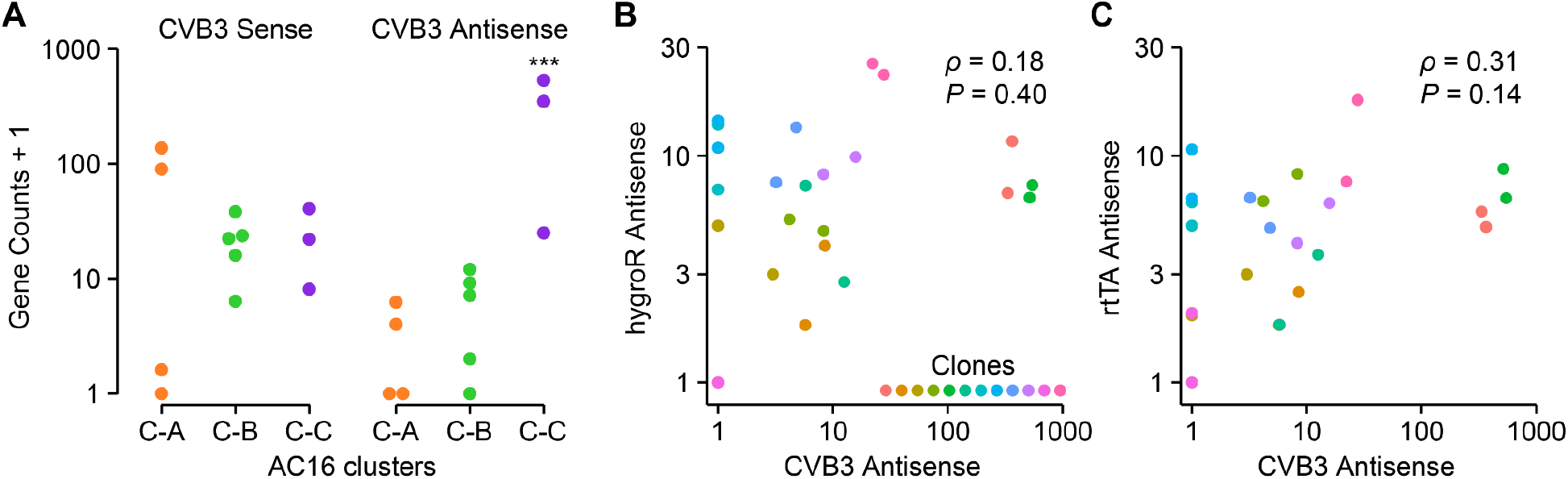
AC16 clone Cluster C (C-C) is distinguished by increased copies of CVB3 antisense genomes. (**A**) Aligned reads of consensus clusters A, B, and C to the sense and antisense genomes of CVB3. \*\*\**P* < 0.001 to C-A or C-B. (**B** and **C**) CVB3 antisense reads are not significantly coupled to other antisense gene cassettes in the linearized CVB3 A67G plasmid transfected (fig. S5).

**Fig. S8.**
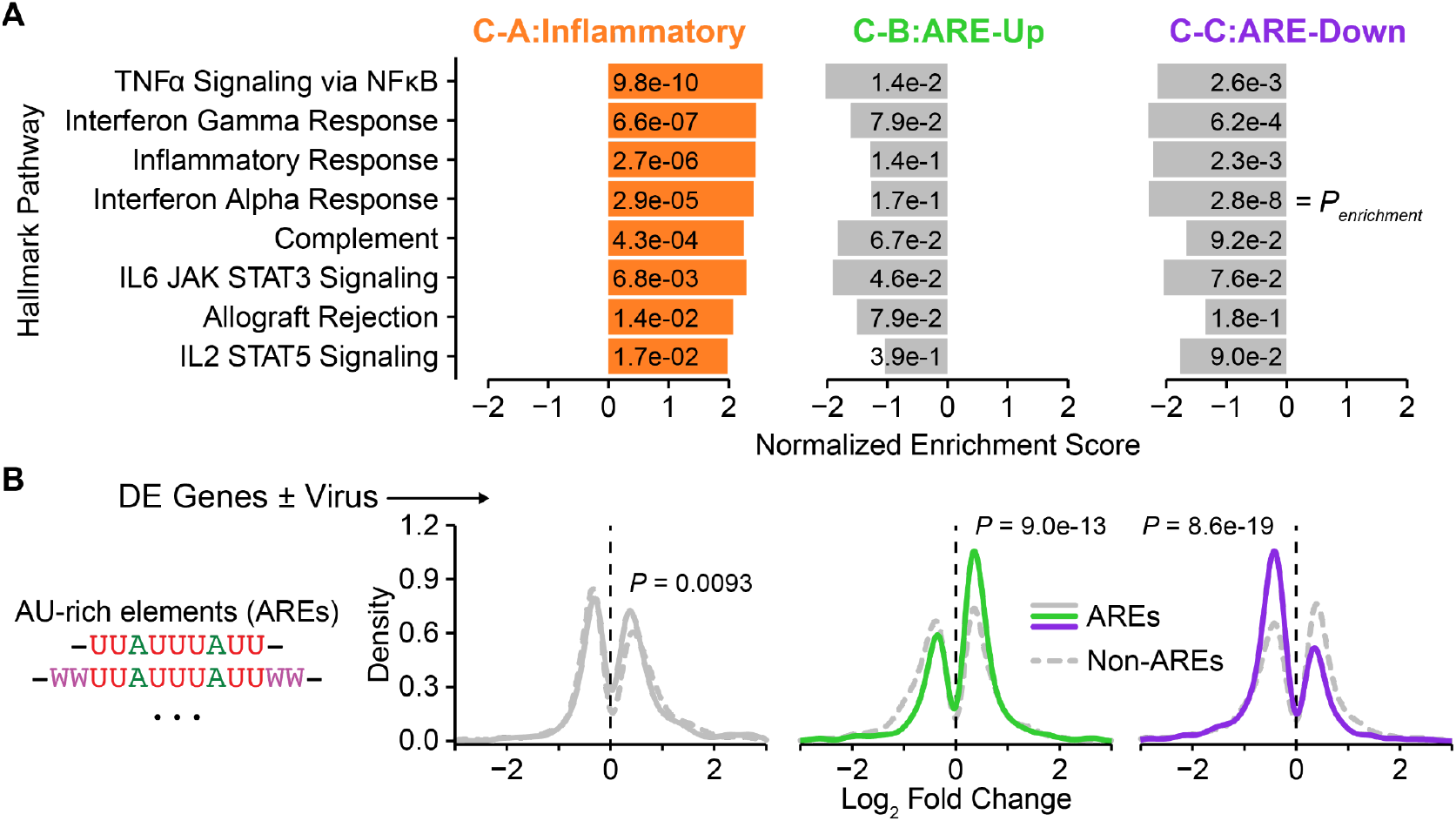
Coxsackievirus B3 adaptions in AC16 cardiomyocytes. (**A**) MSigDB analysis of hallmark pathway enrichments for the three consensus clusters: C-A, Inflammatory; C-B, ARE-Up; and C-C, ARE-Down. The normalized enrichment score is shown for differentially increased (positive) or decreased (negative) genes relative to virus-negative controls. The false discovery rate-corrected *P* value for the enrichment (*P_enrichment_*) is inset. The full set of enrichments is listed in table S9. (**B**) The differentially expressed (DE) genes of C-B and C-C show biased proportions of transcripts with AU-rich elements (AREs). The distributions of ARE-containing genes from the ARED-Plus (*129*) database were plotted along with non-ARE DE genes for each cluster, and the DE genes were assessed for ARE enrichment by the hypergeometric test with all genes (DE and non-DE) as the reference and Bonferroni correction for multiple-hypothesis testing.

**Fig. S9.**
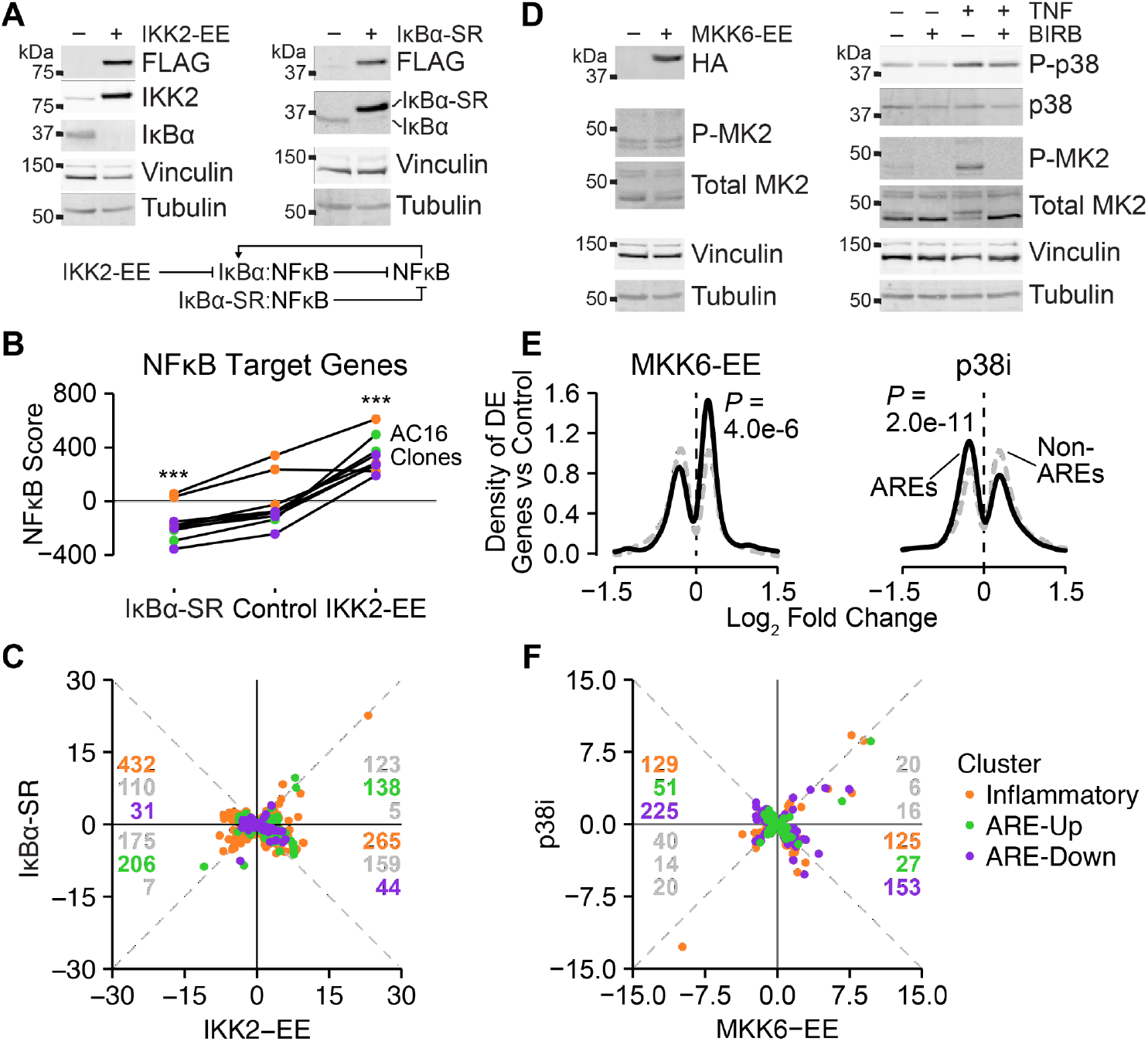
Validation of ARE and inflammatory perturbations. (**A**) Validation of constitutively active IKK2-EE and dominant-negative IκBα super-repressor (IκBα-SR). AC16 cells were acutely transduced, selected, and immunoblotted for FLAG-IKK2, FLAG-IκBα-SR, or IκBα with vinculin and tubulin used as loading controls. The regulatory architecture of IKK2-EE and IκBα-SR on endogenous IκBα and NFκB is shown underneath. (**B**) Projection of CVB3 A67G-engineered AC16 clones (*n* = 9, colored by adaption) on a 200-gene signature of NFκB target genes (*128*) to yield an NFκB score. Differences between the IKK2-EE and IκBα-SR transduced clones and the pBabe control were assessed by paired *t* test. \*\*\**P* < 0.001. (**C**) Log_2_ fold-change correlations between IKK2-EE and IκBα-SR differentially expressed genes in CVB3-expressing AC16 clones (*n* = 9). (**D**) Validation of constitutively active MKK6-EE and the p38 inhibitor (p38i) BIRB796. AC16 cells were acutely transduced–selected or inhibited with p38i (5 µM) for 1 hour before treatment with 20 ng/ml TNF for 30 minutes. Samples were immunoblotted for 3xHA-MKK6-EE, phospho-MK2 (P-MK2), total MK2, phospho-p38 (P-p38), and total p38 with vinculin and tubulin used as loading controls. (**E**) The differentially expressed (DE) genes of MKK6-EE- and p38i-perturbed cells show biased proportions of transcripts with AU-rich elements (AREs). The distributions of ARE-containing genes from the ARED-Plus (*129*) database were plotted along with non-ARE DE genes for all samples, and the DE genes were assessed for ARE enrichment by the hypergeometric test with all genes (DE and non-DE) as the reference. (**F**) Log_2_ fold-change correlations between MKK6-EE and p38i differentially expressed genes in CVB3-expressing AC16 clones (*n* = 9). For (C) and (F), enriched directional correlations were assessed by the hypergeometric test. Colored gene counts indicate Bonferroni-corrected *P* < 0.05 for each adaption.

**Fig. S10.**
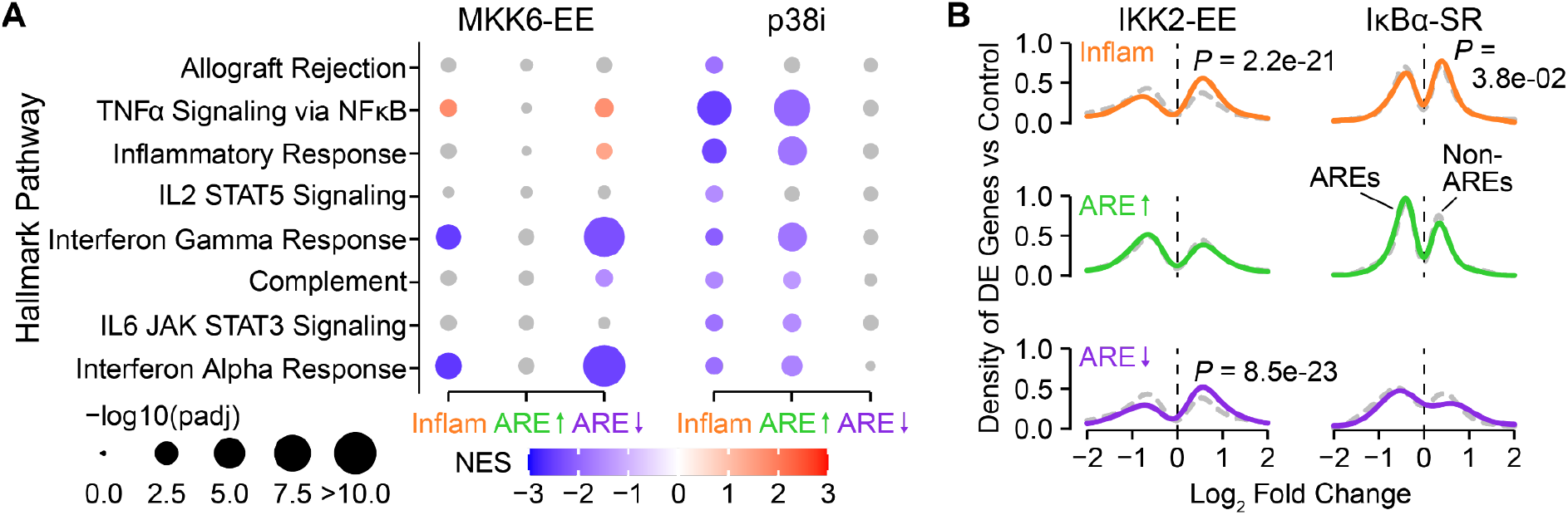
Limited collateral effects of ARE and inflammatory perturbations. (**A**) Differential changes in inflammatory hallmarks for AC16 clones engineered with constitutively active MKK6-EE or treated with the p38 inhibitor (p38i) BIRB796 (5 µM). The normalized enrichment score (NES) was calculated for each inflammatory hallmark pathway relative to empty vector control. The hallmark pathways are ordered as in Fig. 1C. (**B**) Alteration of ARE-Down and ARE-Up states in AC16 clones engineered with constitutively active IKK-EE or dominant-negative IκBα super-repressor (IκBα-SR). The distribution of differentially expressed genes with or without AREs for each adaption was compared as in Fig. 1D.

**Fig. S11.**
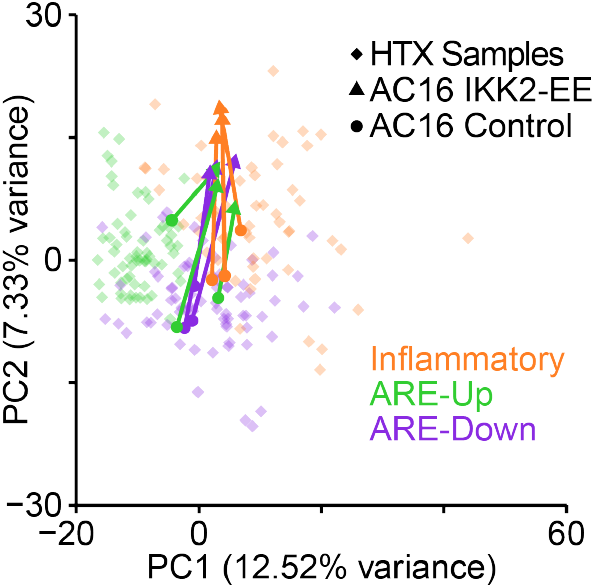
IKK2-EE displacements of AC16 clones within the range of the Inflammatory adaption for HTX samples. Principal component (PC) projection of the differentially expressed genes shared by AC16 clones and HTX samples from Fig. 3G overlaid with AC16 clones engineered with IKK2-EE (triangles) or empty vector control (circles).

**Fig. S12.**
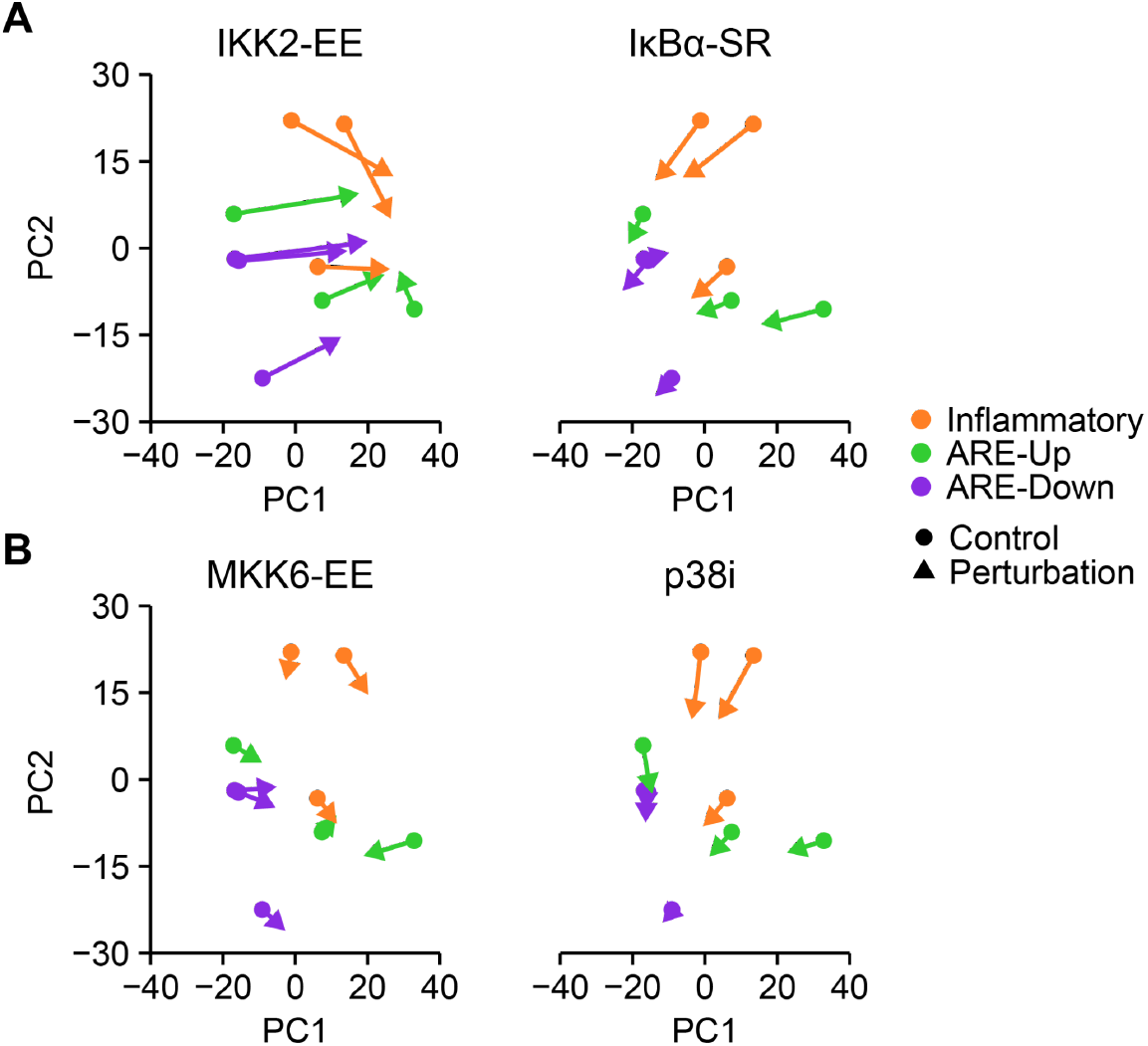
Relocalization of individual AC16 clones upon genetic or pharmacologic perturbation. Principal component (PC) projection of individual AC16 clones within each adaption after the indicated perturbation (triangles) compared to control (circles).

**Fig. S13.**
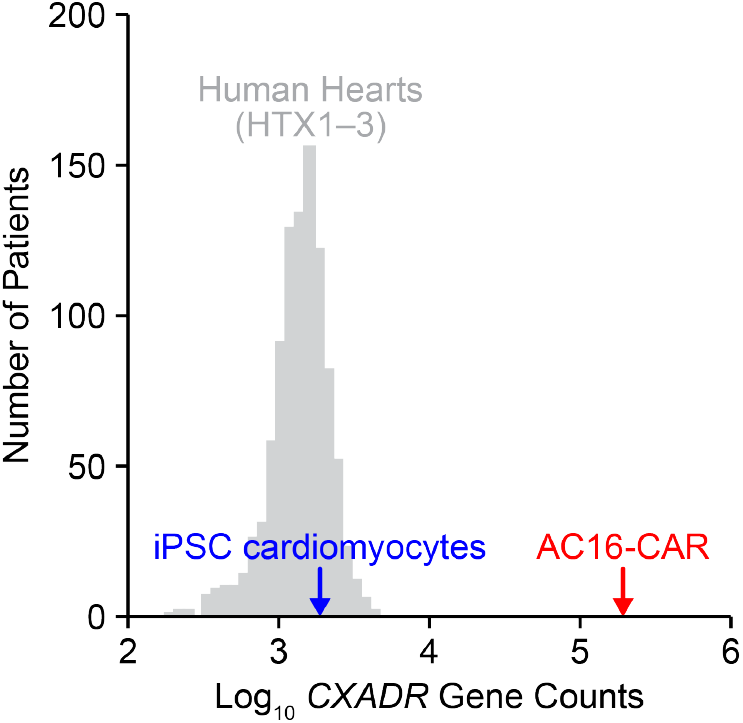
Engineered AC16-CAR cells express supraphysiological copies of CVB3 receptors. Abundance of the coxsackieviral receptor *CXADR* in induced pluripotent stem cell (iPSC)-derived cardiomyocytes and AC16-CAR cells relative to human heart transcriptomes.

**Fig. S14.**
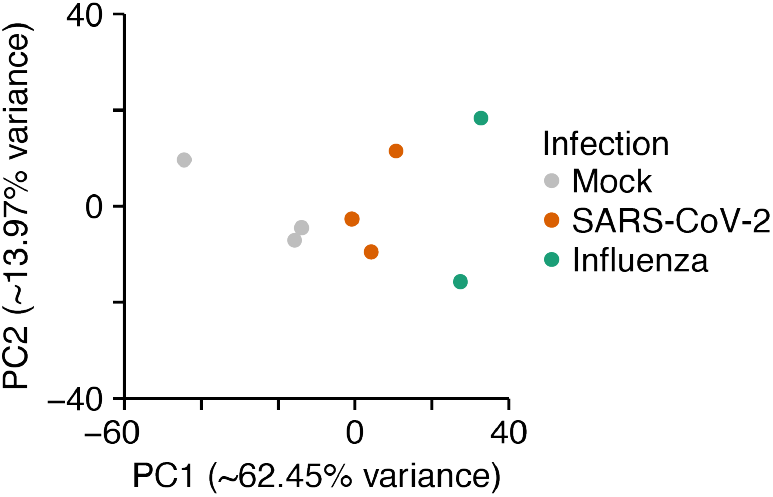
Similar directional changes in gene expression of the heart upon recovery from cardio-pathogen infection. Principal component (PC) projection of the 1000 most-variable genes in hamsters at 31 days after infection with SARS-CoV-2 or influenza A (*67*). PC1 captures ∼4.5x more variance than PC2.

**Fig. S15.**
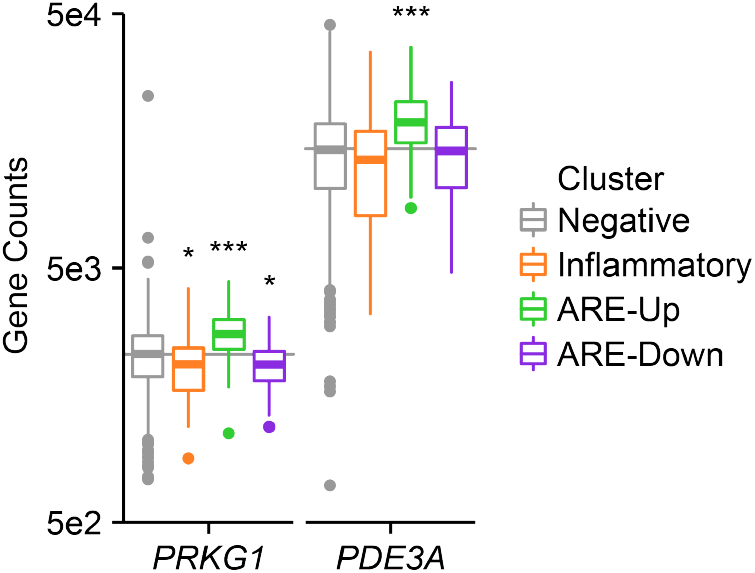
Heart failure drug targets are increased in the ARE-Up adaption. PRKG1 is activated by cGMP produced by guanylate cyclase, which is activated by vericiguat (*135*). PDE3A is inhibited by milrinone (*136*). Abundance of *PRKG1* and *PDE3A* transcripts is separated by adaption, with the virus-negative sample median shown as a horizontal line. Differential gene expression in each adaption was compared to virus-negative samples by DESeq2. \**P* < 0.05, \*\*\**P* < 0.001.

**Table S1.**
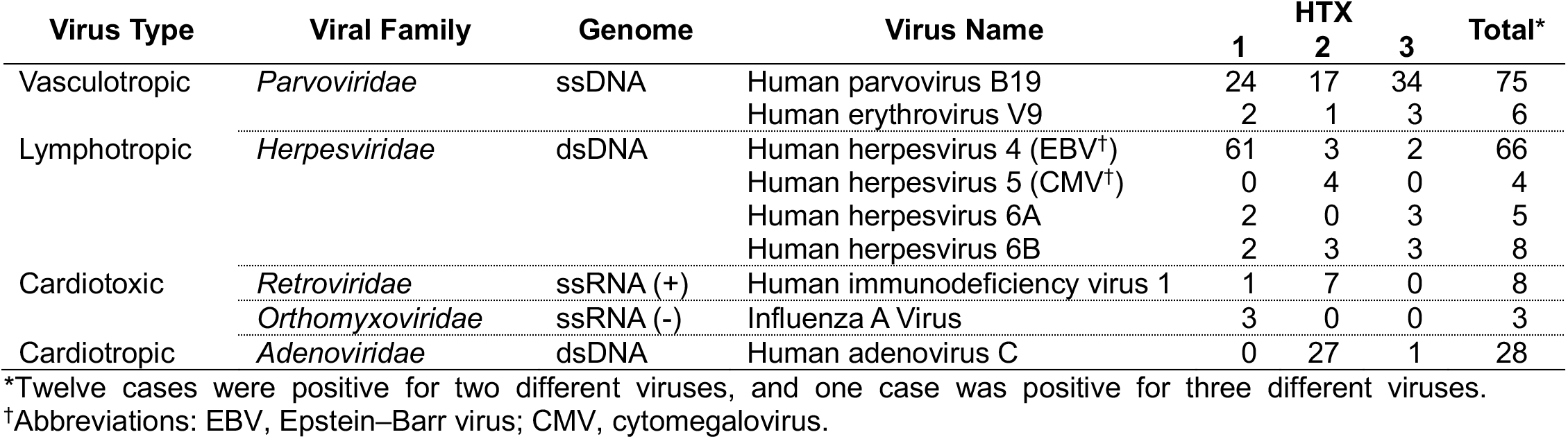
Prevalence of cardio-pathogenic viruses among heart transcriptomics (HTX) datasets.

| Virus Type | Viral Family | Genome | Virus Name | HTX |  |  | Total* |
| --- | --- | --- | --- | --- | --- | --- | --- |
|  |  |  |  | 1 | 2 | 3 |  |
| Vasculotropic | *Parvoviridae* | ssDNA | Human parvovirus B19 | 24 | 17 | 34 | 75 |
|  |  |  | Human erythrovirus V9 | 2 | 1 | 3 | 6 |
| Lymphotropic | *Herpesviridae* | dsDNA | Human herpesvirus 4 (EBV<sup>†</sup>) | 61 | 3 | 2 | 66 |
|  |  |  | Human herpesvirus 5 (CMV<sup>†</sup>) | 0 | 4 | 0 | 4 |
|  |  |  | Human herpesvirus 6A | 2 | 0 | 3 | 5 |
|  |  |  | Human herpesvirus 6B | 2 | 3 | 3 | 8 |
| Cardiotoxic | *Retroviridae* | ssRNA (+) | Human immunodeficiency virus 1 | 1 | 7 | 0 | 8 |
|  | *Orthomyxoviridae* | ssRNA (-) | Influenza A Virus | 3 | 0 | 0 | 3 |
| Cardiotropic | *Adenoviridae* | dsDNA | Human adenovirus C | 0 | 27 | 1 | 28 |

\*Twelve cases were positive for two different viruses, and one case was positive for three different viruses.
<sup>†</sup>Abbreviations: EBV, Epstein–Barr virus; CMV, cytomegalovirus.

**table S2. Universal sample names for HTX datasets.** Each sample is annotated by HTX identifier, disease state, and viral adaption (if applicable). The table also includes the original naming convention, sex, class of cardio-pathogen (if applicable), inferred ancestry, estimated RNA integrity, and total reads in millions.

**table S3. Differentially expressed (DE) genes in virus-positive human heart samples.** Genes are sorted by their FDR-adjusted *P* value (p_adj_ < 0.1) for the (**A**) Inflammatory, (**B**) ARE-Up, and (**C**) ARE-Down clusters. The table also includes the base mean abundance in DESeq2 normalized counts, the log_2_ fold change relative to virus-negative human heart samples, and the standard error (lfcSE).

**table S4. Gene set enrichment analysis (GSEA) of ranked genes in virus-positive human heart samples.** Input genes are ranked by shrunken log_2_ fold change relative to virus-negative human heart samples. Hallmark pathways are sorted by their FDR-adjusted *P* value (p_adj_) for the (**A**) Inflammatory, (**B**) ARE-Up, and (**C**) ARE-Down clusters. The table also includes the normalized enrichment score (NES) for upregulated (NES > 0) and downregulated (NES < 0) hallmarks.

**table S5. Gene signature matrix for bulk sample deconvolution with CIBERSORTx.** Weighting of the indicated transcripts for deconvolving the indicated cell types is shown.

**table S6. Comparisons of AC16 cells overexpressing the CVB3 receptor CXADR (AC16-CAR) infected with wildtype CVB3 or CVB3 A67G.** DE genes in AC16-CAR cells infected with (**A**) wildtype CVB3, (**B**) CVB3 A67G, or (**C**) both. The table also includes the base mean abundance in DESeq2 normalized counts, the log_2_ fold change relative to mock-infected controls, and the standard error (lfcSE). (**D**) GSEA of shrunken log_2_ fold change ranked genes identified in (C) sorted by their FDR-adjusted *P* value (p_adj_). The table also includes the normalized enrichment score (NES) for upregulated (NES > 0) and downregulated (NES < 0) hallmarks.

**table S7. Comparisons of induced pluripotent stem cell-derived cardiomyocytes (iPSC-CM) infected with wildtype CVB3 or CVB3 A67G.** DE genes in iPSC-CM cells infected with (**A**) wildtype CVB3, (**B**) CVB3 A67G, or (**C**) both. The table also includes the base mean abundance in DESeq2 normalized counts, the log_2_ fold change relative to mock-infected controls, and the standard error (lfcSE). (**D**) GSEA of shrunken log_2_ fold change ranked genes identified in (C) sorted by their FDR-adjusted *P* value (p_adj_). The table also includes the normalized enrichment score (NES) for upregulated (NES > 0) and downregulated (NES < 0) hallmarks.

**table S8. DE genes in AC16 clones engineered with CVB3 A67G.** Genes are sorted by their FDR-adjusted *P* value (p_adj_ < 0.1) for the (**A**) Inflammatory, (**B**) ARE-Up, and (**C**) ARE-Down clusters. The table also includes the base mean abundance in DESeq2 normalized counts, the log_2_ fold change relative to the other two AC16 clusters, and the standard error (lfcSE).

**table S9. GSEA of ranked genes in AC16 clones engineered with CVB3 A67G.** Input genes are ranked by shrunken log_2_ fold change relative to the other two AC16 clusters. Hallmark pathways are sorted by their FDR-adjusted *P* value (p_adj_) for the (**A**) Inflammatory, (**B**) ARE-Up, and (**C**) ARE-Down clusters. The table also includes the normalized enrichment score (NES) for upregulated (NES > 0) and downregulated (NES < 0) hallmarks.

**table S10. DE genes in CVB3-expressing AC16 clones transduced with IKK2-EE.** Genes are sorted by their FDR-adjusted *P* value (p_adj_ < 0.1) for (**A**) all clones or clones in the (**B**) Inflammatory, (**C**) ARE-Up, or (**D**) ARE-Down clusters. The table also includes the base mean abundance in DESeq2 normalized counts, the log_2_ fold change relative to pBabe puro + DMSO control, and the standard error (lfcSE).

**table S11. DE genes in CVB3-expressing AC16 clones transduced with IκBα super-repressor (IκBα-SR).** Genes are sorted by their FDR-adjusted *P* value (p_adj_ < 0.1) for (**A**) all clones or clones in the (**B**) Inflammatory, (**C**) ARE-Up, or (**D**) ARE-Down clusters. The table also includes the base mean abundance in DESeq2 normalized counts, the log_2_ fold change relative to pBabe puro + DMSO control, and the standard error (lfcSE).

**table S12. GSEA of ranked genes in CVB3-expressing AC16 clones transduced with IKK2-EE.** Input genes are ranked by shrunken log_2_ fold change relative to pBabe puro + DMSO control. Hallmark pathways are sorted by their FDR-adjusted *P* value (p_adj_) for the (**A**) Inflammatory, (**B**) ARE-Up, and (**C**) ARE-Down clusters. The table also includes the normalized enrichment score (NES) for upregulated (NES > 0) and downregulated (NES < 0) hallmarks.

**table S13. GSEA of ranked genes in CVB3-expressing AC16 clones transduced with IκBα-SR.** Input genes are ranked by shrunken log_2_ fold change relative to pBabe puro + DMSO control. Hallmark pathways are sorted by their FDR-adjusted *P* value (p_adj_) for the (**A**) Inflammatory, (**B**) ARE-Up, and (**C**) ARE-Down clusters. The table also includes the normalized enrichment score (NES) for upregulated (NES > 0) and downregulated (NES < 0) hallmarks.

**table S14. DE genes in CVB3-expressing AC16 clones transduced with MKK6-EE.** Genes are sorted by their FDR-adjusted *P* value (p_adj_ < 0.1) for (**A**) all clones or clones in the (**B**) Inflammatory, (**C**) ARE-Up, or (**D**) ARE-Down clusters. The table also includes the base mean abundance in DESeq2 normalized counts, the log_2_ fold change relative to pBabe puro + DMSO control, and the standard error (lfcSE).

**table S15. DE genes in CVB3-expressing AC16 clones treated with BIRB796 (p38i).** Genes are sorted by their FDR-adjusted *P* value (p_adj_ < 0.1) for (**A**) all clones or clones in the (**B**) Inflammatory, (**C**) ARE-Up, or (**D**) ARE-Down clusters. The table also includes the base mean abundance in DESeq2 normalized counts, the log_2_ fold change relative to pBabe puro + DMSO control, and the standard error (lfcSE).

**table S16. GSEA of ranked genes in CVB3-expressing AC16 clones transduced with MKK6-EE.** Input genes are ranked by shrunken log_2_ fold change relative to pBabe puro + DMSO control. Hallmark pathways are sorted by their FDR-adjusted *P* value (p_adj_) for the (**A**) Inflammatory, (**B**) ARE-Up, and (**C**) ARE-Down clusters. The table also includes the normalized enrichment score (NES) for upregulated (NES > 0) and downregulated (NES < 0) hallmarks.

**table S17. GSEA of ranked genes in CVB3-expressing AC16 clones treated with p38i.** Input genes are ranked by shrunken log_2_ fold change relative to pBabe puro + DMSO control. Hallmark pathways are sorted by their FDR-adjusted *P* value (p_adj_) for the (**A**) Inflammatory, (**B**) ARE-Up, and (**C**) ARE-Down clusters. The table also includes the normalized enrichment score (NES) for upregulated (NES > 0) and downregulated (NES < 0) hallmarks.

**table S18. Comparisons of human hearts infected with SARS-CoV-2.** (**A**) DE genes in human hearts infected with SARS-CoV-2. Genes are sorted by their FDR-adjusted P value (p_adj_ < 0.1). The table also includes the base mean abundance in DESeq2 normalized counts, the log_2_ fold change relative to uninfected controls, and the standard error (lfcSE). (**B**) GSEA of shrunken log_2_ fold change ranked genes in human hearts infected with SARS-CoV-2. Hallmark pathways are sorted by their FDR-adjusted *P* value (p_adj_). The table also includes the normalized enrichment score (NES) for upregulated (NES > 0) and downregulated (NES < 0) hallmarks.

**table S19. Comparisons of hamster hearts infected with SARS-CoV-2 or influenza A.** (**A**) DE genes in hamster hearts infected with SARS-CoV-2 or influenza A. Genes are sorted by their FDR-adjusted P value (p_adj_ < 0.1). The table also includes the base mean abundance in DESeq2 normalized counts, the log_2_ fold change relative to uninfected controls, and the standard error (lfcSE). (**B**) GSEA of shrunken log_2_ fold change ranked genes in hamster hearts infected with SARS-CoV-2 or influenza A. Hallmark pathways are sorted by their FDR-adjusted *P* value (p_adj_). The table also includes the normalized enrichment score (NES) for upregulated (NES > 0) and downregulated (NES < 0) hallmarks.

**Table S20.**
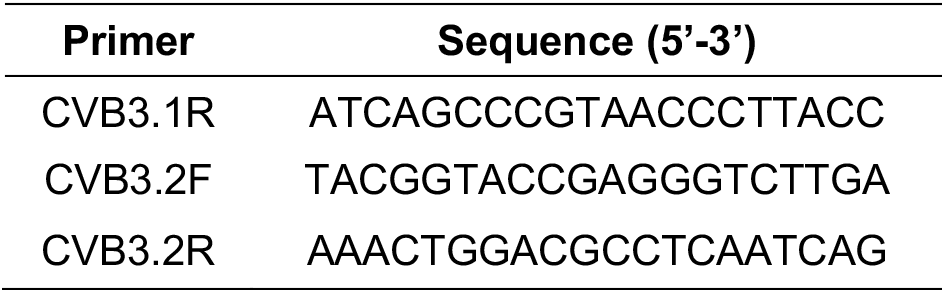
Primer sequences used for CVB3 strand specific quantitative PCR.

